# A gut commensal niche regulates stable association of a multispecies microbiota

**DOI:** 10.1101/2021.09.30.462663

**Authors:** Ren Dodge, Eric W. Jones, Haolong Zhu, Benjamin Obadia, Daniel J. Martinez, Chenhui Wang, Andrés Aranda-Díaz, Kevin Aumiller, Zhexian Liu, Marco Voltolini, Eoin L. Brodie, Kerwyn Casey Huang, Jean M. Carlson, David A. Sivak, Allan C. Spradling, William B. Ludington

## Abstract

The intestines of animals are typically colonized by a complex, relatively stable microbiota that influences health and fitness, but the underlying mechanisms of colonization remain poorly understood. As a typical animal, the fruit fly, *Drosophila* melanogaster, is associated with a consistent set of commensal bacterial species, yet the reason for this consistency is unknown. Here, we use gnotobiotic flies, microscopy, and microbial pulse-chase protocols to show that a commensal niche exists within the proventriculus region of the *Drosophila* foregut that selectively binds bacteria with exquisite strain-level specificity. Primary colonizers saturate the niche and exclude secondary colonizers of the same strain, but initial colonization by *Lactobacillus* physically remodels the niche to favor secondary colonization by *Acetobacter*. Our results provide a mechanistic framework for understanding the establishment and stability of an intestinal microbiome.

**One-Sentence Summary:** A strain-specific set of bacteria inhabits a defined spatial region of the *Drosophila* gut that forms a commensal niche.

## Main Text

Animal guts are colonized by a complex community of host-specific commensal bacteria that is relatively stable over time within an individual (*1–3*) and can have life-long effects on health (*4, 5*). It is unknown how this microbiome is established and maintained over time in the face of daily fluctuations in diet (*6*), invasion by pathogens (*7*), and disruptions by antibiotics (*8*). One hypothesis is that long-term maintenance of diet and lifestyle habits reinforces microbiome stability (*1, 9*), while an alternative, non-exclusive hypothesis is that the host constructs microbial niches in the gut that acquire and sequester symbiotic bacteria (*10–14*).

The microbiome of the fruit fly, *Drosophila melanogaster*, has been studied for over a century and is relatively simple in its composition compared to the mammalian gut (*15*), yet how gut microbiome assembly is regulated remains unclear. Like human colonic crypts, the fly gut is microaerobic and colonized by bacteria from the Lactobacillales class and Proteobacteria phylum (*16–19*). Flies can easily be reared germ-free and then associated with defined bacterial strains, providing a high level of biological control (*20*). Furthermore, the fly gut microbiota are of low diversity, with ∼5 species of stable colonizers from two primary groups: the genera *Lactobacillus* (phylum Firmicutes), which was recently split into *Lactiplantibacillus* and *Levilactibacillus,* and *Acetobacter* (class *α*-Proteobacteria) (*19, 21*). These species are easily cultured, genetically tractable (*20*), and they affect fly lifespan, fecundity, and development (*22–28*).

While colonization of the fly gut has long been argued to be non-specifically regulated by host filtering mechanisms, including feeding preferences, immunity, and digestion, recent evidence suggests flies may also selectively acquire *Lactobacillus* and *Acetobacter* strains in the wild (*17, 29*). Here, we discover an ecological niche within the *Drosophila* foregut and characterize priority effects that regulate the stable gut association of specific bacterial species.

## Results

### Spatially specific gut localization of Lactobacillus plantarum from wild flies

To investigate whether commensal bacteria form stable associations with the fly gut in a manner consistent with the existence of a niche, we exposed flies to a quantified inoculum of bacterial cells labeled with a fluorescent protein (Fig. S1A-G). Following inoculation, flies were transferred to germ-free food daily for 3 d followed by an additional transfer to a new germ-free vial for 3 h to allow transient bacteria to clear from the gut (Methods, Fig. S1). Clearing prior to analysis reduced the total number of gut bacteria and the spatial variation in bacterial location (Fig. S1H-J). These experiments revealed that a strain of *Lactobacillus plantarum* (*Lp*) isolated from a wild-caught fly (*LpWF*) persists exclusively in the *D. melanogaster* foregut (Fig. 1A-E, S1I,J), including the proventriculus (a luminal region connecting the esophagus with the anterior midgut (*30*)), the crop (a sack-like appendage), and the crop duct that connects the crop to the proventriculus. Bacteria associated with longitudinal furrows lining the surface of the proventriculus inner lumen, the crop duct, and the base of the crop (Fig. 1C-E, S1J). Similar to *LpWF*, a strain of *Acetobacter indonesiensis* colonized the same foregut regions (Fig. 1F, S2), indicating that the two major groups of fly gut bacteria have the same spatial specificity in the foregut. By contrast, flies colonized with *Lp* from laboratory flies (*LpLF*) (Fig. S1K) or the *LpWCFS1* strain isolated from humans (Fig. S1L) had much lower levels of colonization. No *Lp* strains were found at substantial abundance in the midgut or other regions of the fly after clearing transient bacteria. Consistent with microscopy, live bacterial density was greatest in the proventriculus, followed by the crop, and was lowest in the midgut and hindgut (Fig. 1G, S1M). We further validated that *LpWF* maintains stable colonization in the absence of ingestion of new bacterial cells over 5 d during which non-adherent bacteria were flushed from the gut by fastidiously maintaining sterility of the food using a CAFÉ feeder (*17*) (Fig. S3A,B).

**Fig. 1.**
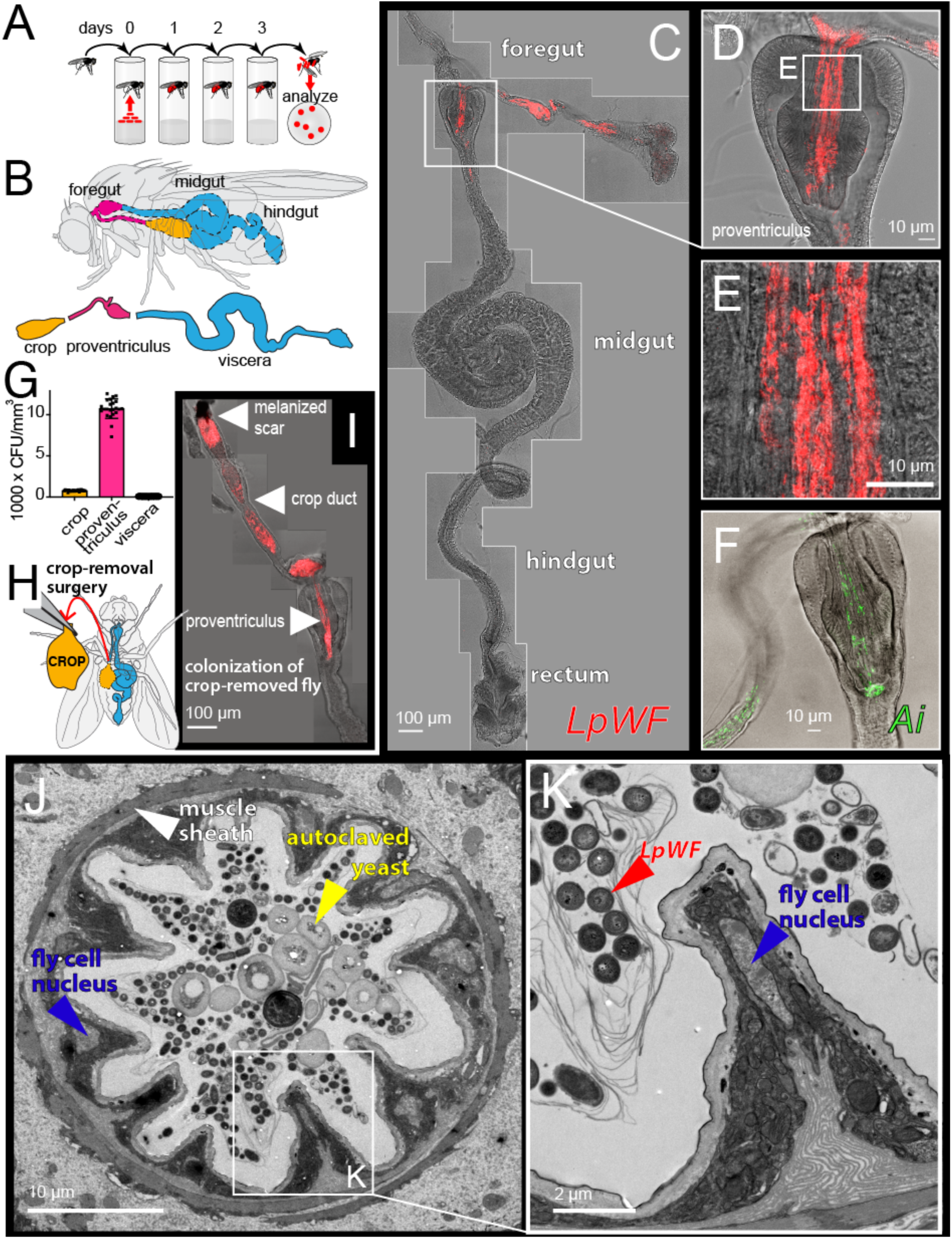
*LpWF* stably colonizes the fly gut with spatial specificity. (**A**) Colonization assay schematic. (**B**) Gut diagram. (**C**) Microscopy of *LpWF*-mCherry colonization in whole gut after clearing transient cells shows a specific colonization zone in the foregut. Max intensity z-projection. Scale bar: 100µm. (**D**) Proventriculus. (**E**) Anterior proventriculus inner lumen. (**F**) *Ai* colonization is also specific to the proventriculus lumen and crop duct. (**G**) CFU densities from regions dissected in B. *n* = 60 individual guts/region. (**H**) Microsurgery to remove the crop. (**I**) *LpWF* colonizes the foregut of flies with the crop removed (*n*=15/15). Arrow 1: healed wound site. Arrow 2: crop duct. Arrow 3: proventriculus (c.f. panel C). (**J**) TEM cross section of proventriculus inner lumen. (**K**) Detail of J.

A bacterial population in the foregut with the observed spatial localization might be maintained by proliferation and constant re-seeding from the crop, in which case flies without crops could not be stably colonized. We conducted microsurgery to remove the crop from germ-free flies (Fig. 1H, Methods), inoculated them with *LpWF* 5 d post-surgery, and then dissected and imaged the gut 5 d post inoculation (dpi). Surgical removal was validated and the remaining portion of the crop duct had a melanized scar at the surgery site (Fig. 1I). All cropless flies were stably colonized by *LpWF* (*n*=15/15), with a high density of bacteria in the proventriculus inner lumen as in flies with an intact crop (c.f. Fig. 1C). We observed similar *Ai* colonization following cropectomy (n=14/14 colonized; Fig. S2D,E). Thus, the crop is not required for stable foregut colonization by *LpWF* or *Ai*, suggesting that the ability of the bacteria to specifically bind to the proventriculus and crop duct is key to stable bacterial association. Examining these regions further, transmission electron microscopy (TEM) of the proventriculus lumen revealed a consistent tissue geometry (Fig. 1J,K), with densely packed bacterial cells longitudinally oriented in elongated furrows formed by host cell bodies making up an average of 11 ridges per cross section (Fig. S3C).

### Commensal association saturates at a precise bacterial population size and resists displacement, suggesting a niche

A niche would be expected to result in strong bacterial association based on specific binding sites, such that the associated bacterial population size would saturate at a well-defined value. Moreover, cells already bound to the proventriculus would be expected to promote population stability and prevent later-arriving bacteria from colonizing. To test these hypotheses, we colonized germ-free flies with a range of doses of *LpWF*-mCherry and measured the abundance over time. As predicted, over a wide range of initial inoculum sizes, the associated bacterial population saturated at ∼10^4^ CFUs/fly (Fig. 2A). Furthermore, when the inoculum size was below that saturation level, the population of bacteria in the proventriculus increased gradually and plateaued within 5 d. Growth measurements in live flies (*17*) demonstrated that the plateau was reached by growth of the initially bound population rather than ingestion of additional cells. By contrast, when an excess of bacteria was supplied initially, the population decreased to the same plateau value within 1 d (Fig. 2A), indicating that the niche has a finite and fixed carrying capacity. Similar dynamics were observed for *Ai* with ∼10^3^ cells at the saturated density (Fig. S2F).

**Fig. 2.**
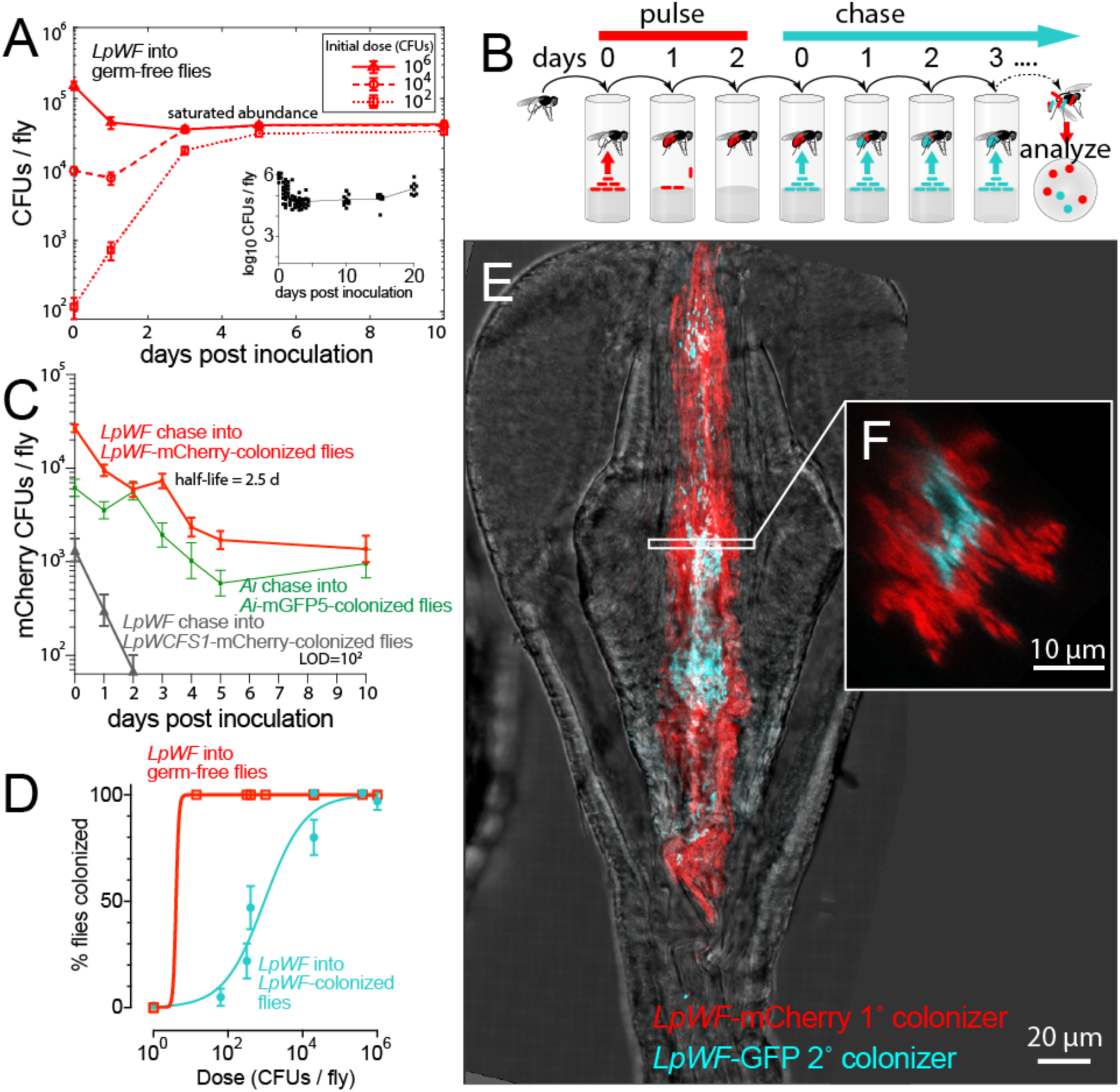
Kinetic properties of bacterial associations suggest the existence of a niche in the proventriculus. (**A**) Saturation occurs in a time course of colonization in germ-free flies inoculated with *LpWF*. Error bars: s.e.m. Inset: 20-day time course after inoculation with 10^6^ CFUs (data from (*17*)). (**B**) Bacterial pulse-chase experimental design: flies were first pre-colonized with *LpWF*-mCherry, then fed an excess of unlabeled *LpWF* (blue) daily on fresh food. (**C**) Bacterial cell turnover quantified by pulse-chase time course of *Lp*-mCherry-pre-colonized flies continuously fed unlabeled *LpWF* or *Ai*-GFP-pre-colonized flies continuously fed unlabeled *Ai.* Error bars: s.e.m. (**D**) Colonization efficiency quantified by dose response to colonization of individual flies. CFUs quantified at 3 dpi of the second colonizer. *n*=24 flies/dose, error bars: standard error of the proportion. Limit of detection: 50 CFUs. (**E**) Spatial structure of colonization dynamics in the proventriculus for a fly pre-colonized with *LpWF*-mCherry (red) invaded by *LpWF*-GFP and imaged 1 hour post inoculation (hpi). (**F**) Optical *x,z-*slice.

To investigate the stability of bacterial colonization in the proventriculus, we performed a pulse-chase experiment in which we challenged *LpWF*-mCherry-pre-colonized flies with unlabeled *LpWF* fed in excess over the course of 10 d (Fig. 2B). *LpWF*-mCherry levels in the gut decreased by >90% over the first 5 d, from ∼10^4^ to ∼10^3^ CFUs/fly, and then remained at ∼10^3^ CFUs/fly for the following 5 d (Fig. 2C), indicating a small, bound population with little turnover and a larger associated population with a half-life of 2.5 d (95% c.i. 1.6 to 4.3 d). By contrast, *LpWCFS1*, a weakly-colonizing human isolate of *L. plantarum*, was quickly flushed from the gut (Fig. 2C). Similar dynamics were observed in *Ai* (Fig. 2C) with a half-life of 2.5 d (95% c.i. 1.3 to 6.5 d), indicating the niche has equivalent kinetic for both bacterial species.

Initial binding to the niche is a key step in the establishment of a new bacterial population prior to filling the niche. Establishment is dose-dependent (*17*), and our finding that the final abundance of late colonizers is lower than that of initial colonizers (Fig. S3D) suggested that the presence of prior colonizers would shift the dose-response curve. To quantify such priority effects, we fed a range of doses of *LpWF*-mCherry to individual *LpWF*-pre-colonized flies and measured the percentage that were colonized by *LpWF*-mCherry 3 d later. Consistent with our hypothesis, pre-colonized flies were less likely than germ-free flies to become colonized by an equal dose of *LpWF*-mCherry: ∼10^3^ *LpWF*-mCherry CFUs were required for 50% of flies to be colonized, while 100% of germ-free flies ended up colonized by doses as low as 10^2^ CFUs (Fig. 2D). These findings demonstrate that the proventricular niche for *LpWF*, when occupied, strongly resists colonization by later doses of the same strain.

The relationship between the probability of establishment and the final abundance of successful colonizing bacteria suggests that the availability of open habitat regulates the chance of invasion. We formalized assumptions of this hypothesis by building an integrated theory of initial colonization (*17*) and niche saturation (*31*) that predicts the likelihood of colonization, 𝑃(𝑁_0_), of an invading species inoculated at a dose of 𝑁_0_ as a function of the final abundance of the invading species, 𝐴(𝑁_0_),

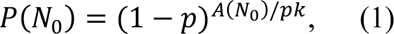

where 𝑝 is the colonization probability of an individual bacterial cell and 𝑘 is the subpopulation size attained in a single successful colonization event Fig. S4A,B). This model allows us to estimate the scale at which the population is structured based on colonization probabilities and total bacterial abundances. For *LpWF,* Eq. 1 estimates a subpopulation size of 𝑘=600 cells (Fig. S4C), which is roughly the number of cells contained in an individual furrow.

To test whether the later dose of *LpWF*-mCherry was spatially excluded by the resident *LpWF*, we constructed a GFP-expressing strain of *LpWF* and fed it to flies pre-colonized with *LpWF*-mCherry. We imaged whole fixed guts 1 h post inoculation (hpi) to capture *LpWF*-GFP cells before they passed out of the fly (Fig. 2E). In the proventriculus, the invading *LpWF*-GFP were localized along the central axis of the inner lumen, separated from the lumen wall by a layer of resident *LpWF*-mCherry (Fig. 2E,F) that was up to 10 µm thick. The posterior proventriculus furrows were densely packed with *LpWF*-mCherry, while *LpWF*-GFP was largely absent from furrows, suggesting that these furrows are the sites of stable colonization. We confirmed that the fluorophores are not responsible for the differential colonization by feeding *LpWF*-mCherry to flies pre-colonized by unlabeled *LpWF* and quantifying the mCherry signal along the gut at 1 hpi and 24 hpi. At 24 hpi with a dose of ∼10^4^ CFUs, flies pre-colonized by *LpWF* showed almost undetectable mCherry by microscopy (Fig. S3E-H). These results provide further support that the niche for *LpWF* is in the proventricular furrows. Unlike during initial colonization, in which bacteria rapidly enter and colonize the furrows, prior colonizers prevent subsequent colonization, suggesting that there are a limited number of binding sites in the furrows for *LpWF* cells and that these sites are saturated by prior colonization. Consistent with this logic that niche priority is spatially-determined, in the cases when *LpWF*-GFP did show colonization (*n*=5), the GFP-labeled cells were co-localized with each other a furrow rather than being evenly mixed with mCherry throughout the proventriculus (Fig. S3I).

### *Ai* and *LpWF* occupy separate niches within the proventriculus

Interspecies interactions can have major impacts on ecosystem colonization through priority effects that include competitive exclusion and facilitation (*32–36*). Because *Ai* and *LpWF* colonize the same general location of the gut (Fig. 1C-F, S2A-C) and each strain excludes itself (Fig. 2D, 3A), we expected that they would exclude each other. To test this hypothesis, we measured each species’ abundance and growth rate during co-colonization. To our surprise, both were unaffected (Fig. 3B,C, S5), demonstrating that the species have independent saturation of the niche. We also performed a dose-response assay to determine whether interactions affect establishment of new colonizers. By contrast to *Ai*’s self-exclusion, *Ai* colonization was facilitated by *LpWF* pre-colonization (Fig. 3A), while *LpWF* colonization was unaffected by the presence of *Ai* (Fig. S5A).

**Fig. 3.**
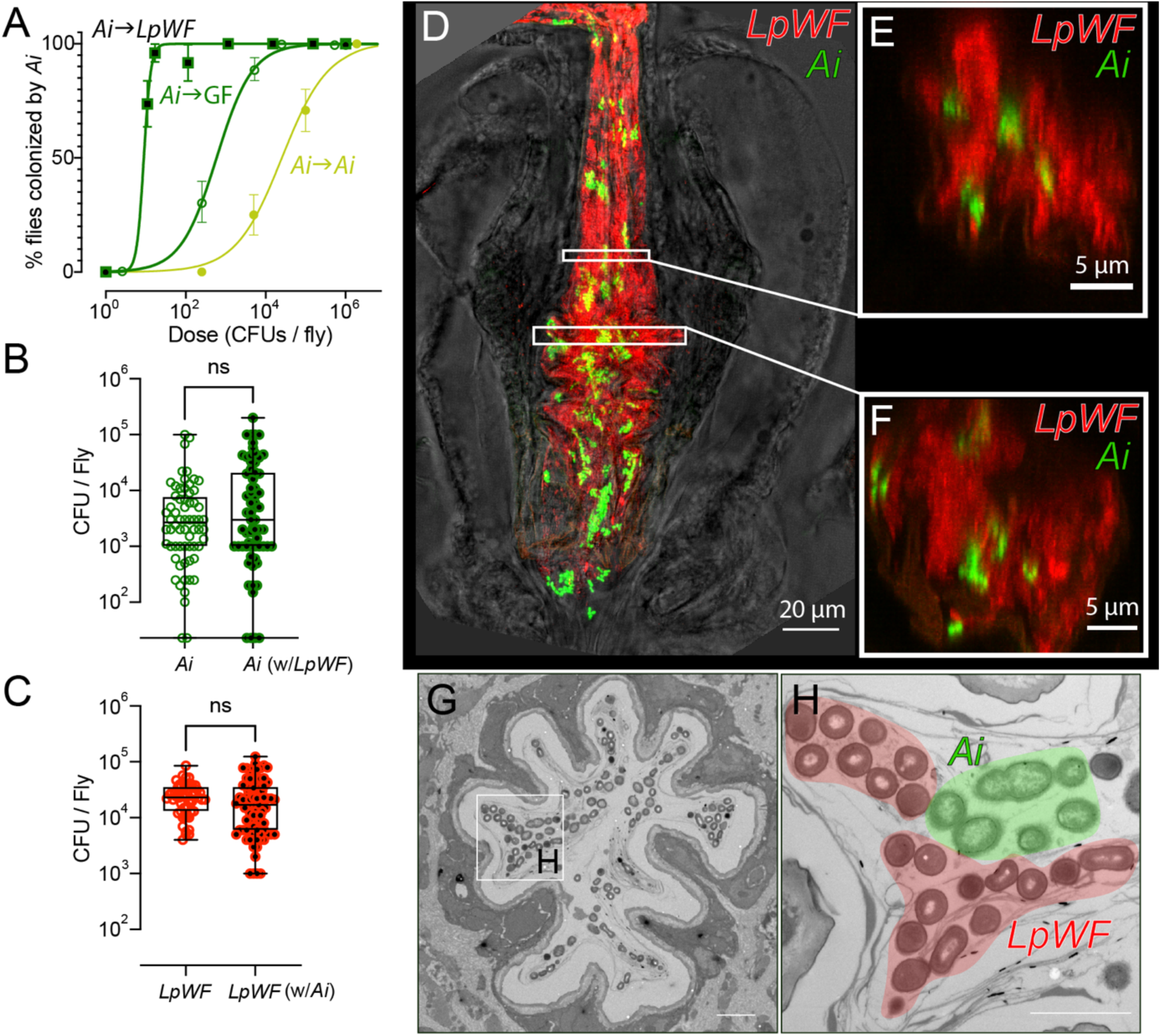
*Ai* and *LpWF* occupy separate niches within the proventriculus. (**A**) Strain interactions influence colonization efficiency as seen by dose-response curve for *Ai* fed to germ-free flies (open green circles), *Ai*-pre-colonized flies (filled yellow circles), or *Lp*-pre-colonized flies (black-filled green squares). Z-test of differences in proportion versus *Ai* into germ-free flies: dose 10^2.3^ CFUs/fly, *p*=8.1×10^- 4^; dose 10^3.7^ CFUs/fly: *p*=4.8×10^-9^; dose 10^5^ CFUs/fly: *p*=8.7×10^-6^). Error bars: standard error of the proportion. (**B**) *Ai* abundance at 5 dpi does not differ between flies monocolonized with *Ai* versus pre-colonized with *LpWF* then fed *Ai*. (**C**) *LpWF* abundance 5 dpi does not differ between flies monocolonized with *LpWF* versus pre-colonized with *Ai* then fed *LpWF* (*n*=60 flies per treatment). (**D**) Confocal microscopy of *Lp* and *Ai* co-colonization. *Ai* (green) and *LpWF* (red) occupy the same regions of the foregut 1 dpi. Scale bar: 100 µm. (**E**,**F**) *x*,*z*-section of *Ai* and *LpWF* sectors. (**G**) TEM cross-section of *Ai* and *LpWF* co-colonized anterior proventriculus. Scale bar: 5 µm. (**H**) Detail of G with *LpWF* and *Ai* cells pseudocolored. Scale bar: 2 µm.

Fluorescence microscopy of guts co-colonized by *LpWF*-mCherry and *Ai*-GFP showed that *Ai* and *LpWF* co-colonized the same foregut regions (Fig. 3D), with distinct sectors of each species observed at the cellular scale (Fig. 3E,F). Thus, *LpWF* and *Ai* do not physically exclude one another, and instead the tissue appears to accommodate both strains.

### Colonization of the niche induces morphological alteration of the proventriculus

To examine the coexistence of overlapping *Ai* and *LpWF* populations in a physically confined space, we imaged fly anatomy using X-ray microcomputed tomography (XR µCT) (*37, 38*), and segmented the volumetric image data to produce 3D reconstructions (Fig. 4A). We imaged germ-free flies and flies colonized with *LpWF*, *Ai*, or both *LpWF* and *Ai*. Numerous crypts were apparent along the length of the gut, including in uncolonized regions of the midgut and hindgut that are shielded by peritrophic matrix (Fig. 4A, S6) (*39*). In the colonized region of the foregut, the longitudinal striations where we observed bacteria coincided with ridges and furrows of host tissue in the proventriculus inner lumen and crop duct (Fig. 4B-F). The furrows were straight in the anterior proventriculus, becoming larger and more irregular in the posterior (Fig. 4D,F). Transverse slices of the lumen wall revealed a narrow passage through the germ-free proventriculus (Fig. 4C), while the opening was much broader in the colonized proventriculus (Fig. 4E), corresponding to a significantly higher luminal volume than in germ-free flies (Fig. 4G).

**Figure 4.**
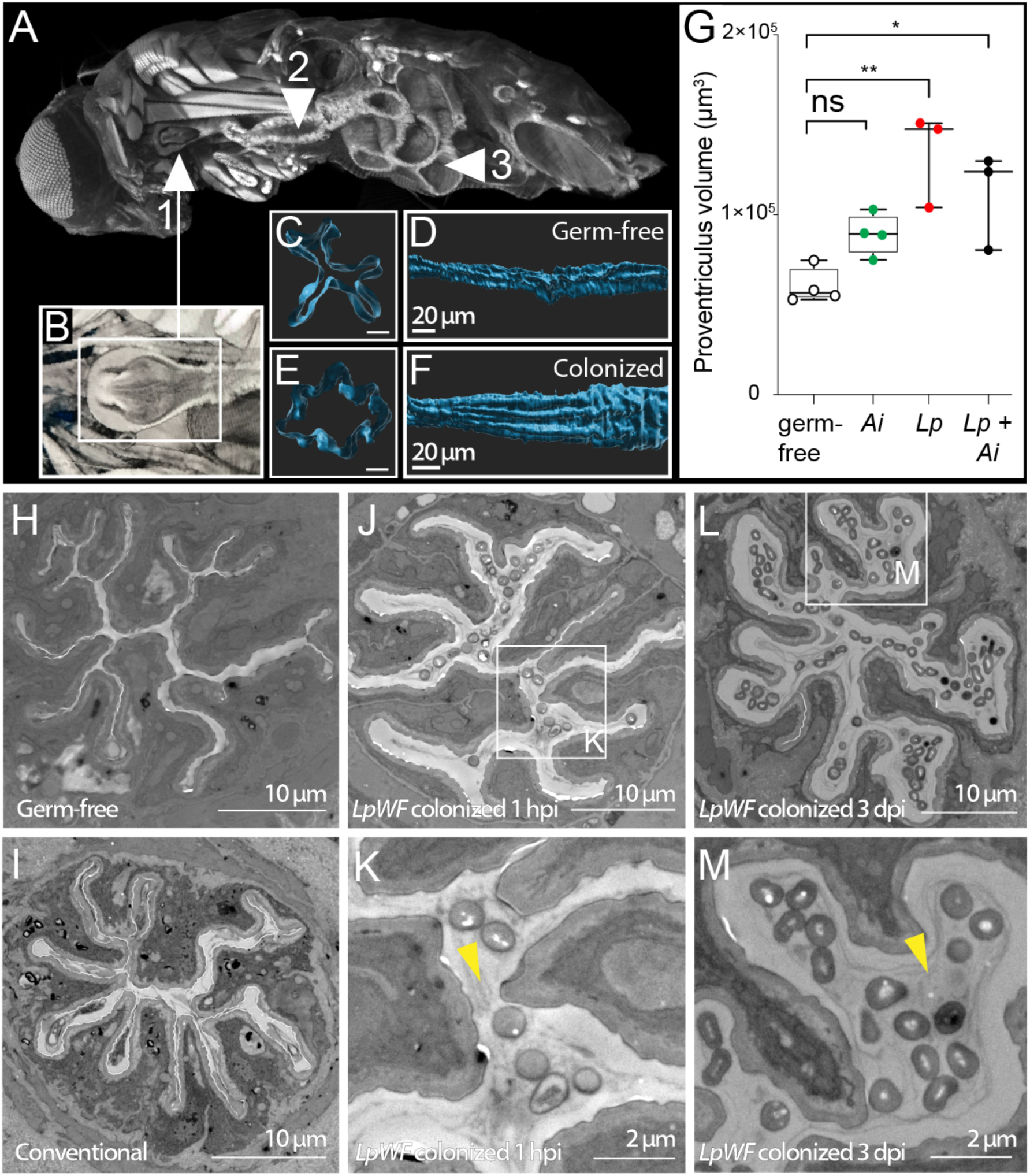
Colonization of the niche induces morphological alteration of the proventriculus. (**A**) XR µCT model of a whole fly. Cutaway shows exposed proventriculus (1, inset B), (2) anterior midgut, and (3) posterior midgut. (**B**) Detail of proventriculus. (**C**) Cross-section of germ-free proventriculus inner lumen. Scale bar: 5 µm. (**D**) Germ-free proventriculus inner lumen volume rendering. Scale bar: 50µm. (**E**) *LpWF*-colonized proventriculus inner lumen cross-section. Scale bar: 10 µm. (**F**) *LpWF* proventriculus inner lumen volume rendering. Scale bar: 50 µm. (**G**) Cardia volume calculated from surface models (*n*=3 to 4 surfaces per condition; *p*=0.0025, one-way ANOVA relative to GF). (**H**) Transmission electron microscopy transverse cross-section of anterior proventriculus in germ-free fly, (**I**) conventionally-reared fly (only lab fly bacteria; no *LpWF*), (**J-K**) 1 hpi with *LpWF*, (**L-M**) 3 dpi colonized with *LpWF* (see Fig. S7), Yellow arrowheads indicate lumen space.

Consistent with XR µCT imaging, TEM cross-sections of the proventriculus of germ-free flies showed a narrow luminal space, approximately 0.5 µm in diameter (Fig. 4H, S7). Similar morphology was observed in conventionally-reared lab flies, which do not have the wild fly strains of bacteria (Fig. 4I). In *LpWF*-colonized flies, the diameter of the furrows increased to ∼1µm by 1 hpi (Fig. 4J,K, S7) and ∼2 to 3 µm by 3 dpi (Fig. 4L,M, S7E-J), suggesting a sustained host response to niche occupancy. The expanded luminal space of the colonized proventriculus contained two zones: a clear zone adjacent to the lumen wall, and a bacteria-colonized zone closer to the center of the lumen (Fig. 4L,M, S7E-J). High pressure freezing fixation (Fig. S7S) suggested that the zonation is not simply an artifact of fixation. This morphology is reminiscent of mammalian mucus, which has two layers: a dense, uncolonized layer adjacent to the epithelium, and a thinner, distal layer colonized by bacteria (*40*). Taken together, our imaging results show that the proventriculus undergoes morphological changes upon colonization, which coincide with the promotion of *Ai* colonization.

## Discussion

Our results show that specific strains of *Drosophila* gut bacteria colonize crypt-like furrows in the proventriculus (Fig. 1C-F, 2E-F, S2, 3D-F, S3I, 4H-M), that the colonization by these strains is saturable (Fig. 2A, 3B,C, S2F), suggesting a limited number of binding sites, and that the proventriculus responds to colonization through engorgement (Fig. 4), which promotes colonization by bacteria that benefit the fly (Fig. 3A) (*25, 26, 41*). The finding that *Drosophila* has a specific niche for binding of commensals to sites in the crop duct and proventriculus is highly significant because it provides insight into how a microbiome can interact with the host in a manner that can be host-regulated and mutually beneficial. Furthermore, it predicts the existence of specific molecules on the surface of the proventriculus that bind to the bacterial surface of colonization-competent strains but not with non-colonizing strains. The finding that binding of one strain can lead to structural changes that open up niche sites for a second species provides a model for how complex assemblies of bacterial strains can arise and be maintained within a host digestive tract.

Despite the long history of studies on the *Drosophila* microbiome, the existence of a specific niche has been obscured by the presence of bacteria in the food and on the culture medium during traditional culturing. A substantial fraction of gut bacteria under such conditions simply pass through and do not interact specifically with the gut (*42*), even though specific microbiome members bound to their associated niches might be present. We used bacterial pulse-chase protocols to push out unbound bacteria, greatly enriching for only specifically interacting cells.

Possession of a microbiome is clearly highly beneficial for *Drosophila*, given that axenic flies show strongly reduced growth and fecundity (*22–25, 43, 44*). However, it is less clear how the relationship between the host and specific strains of bacteria is stably perpetuated. We suggest that understanding the proventricular niche is likely to provide insight into microbiome function 1) by revealing the spatial locations where bacteria influence the host to introduce molecules into the gut, perhaps along with the peritrophic membrane; and 2) by revealing whether changes in niche structure induced by one species lay the groundwork for more complex associations between different members of the microbiome, such as *LpWF* and *Ai*, that are related to their functional pathways. Finally, these observations raise the question of whether additional niches exist at other locations in the *Drosophila* digestive system and within the gut of many other animals, including humans.

## Acknowledgements

Plasmid pCD256-mCherry (*45*) was generously provided by Reingard Grabherr, PhD of BOKU, Austria. Plasmid pCM62 (*46*) was generously provided by Elizabeth Skovran, PhD of SJSU, USA.

## Funding

Banting and Pacific Institute for the Mathematical Sciences Postdoctoral Fellowships (EWJ)

Howard Hughes Medical Institute International Student Research fellow (AA-D)

Stanford Bio-X Bowes fellow (AA-D)

Siebel Scholar (AA-D)

Allen Discovery Center at Stanford on Systems Modeling of Infection (KCH)

Chan Zuckerberg Biohub (KCH)

The David and Lucile Packard Foundation (JMC)

Institute for Collaborative Biotechnologies through Grant W911NF-09-0001 from the US Army Research Office (JMC)

Natural Sciences and Engineering Research Council of Canada Discovery Grant (DAS)

Canada Research Chairs program (DAS)

The Howard Hughes Medical Institute (ACS, CW)

National Institutes of Health grant DP5OD017851 (WBL, BO)

National Science Foundation grant IOS 2032985 (WBL, RD, DM)

Carnegie Institution for Science Endowment (WBL, HZ, KA, RD, DM)

Carnegie Institution of Canada grant (WBL, DAS, EWJ)

National Institutes of Health training grant T32GM007231 (KA, ZL)

## Author Contributions

Research Design: RD, EWJ, HZ, BO, CW, EB, JMC, DS, AS, WBL

Performed Research: RD, EWJ, HZ, BO, DM, CW, KA, AA-D, MV

Analyzed Data: RD, EWJ, WBL

Wrote manuscript: RD, WBL

Revised the Manuscript: RD, EWJ, KCH, DAS, JMC, ACS, WBL

All authors reviewed the manuscript before submission.

## Data and materials availability

All data are available in the main text or the supplementary materials.

## Materials and Methods

### Fly strains and rearing

All flies in this study were mated females, which show low heterogeneity in gut morphology (*45*). Previous work showed that the colonization phenotypes we measure are general across multiple genetic backgrounds including CantonS, *w1118*, and OregonR (*17*). Flies were reared in Wide Drosophila Vials (Cat #: 32-114, Genesee), with Droso-Plugs® (Cat #: 59-201, Genesee). Food composition was 10% glucose (filter-sterilized), 5% autoclaved live yeast, 0.42% propionic acid (filter-sterilized), 1.2% autoclaved agar, and 0.5% cornmeal. Each vial contained 4 mL of food. Germ free fly stocks were passaged to fresh vials every 3-4 d. Five day-old mated female adults were sorted the day prior to beginning an experiment.

Liquid food was composed of 10% glucose, 5% yeast extract, and 0.42% propionic acid. The only nutritional difference between liquid and solid food was yeast extract instead of autoclaved live yeast because the yeast cell walls clog the capillaries used for liquid feeding. The bottom of capillary feeder vials contained 1.2% agar as a hydration and humidity source. Both CAFÉ- and solid food-fed flies were transferred daily to fresh vials to minimize bacterial re-ingestion. Samples of flies were surface-sterilized and crushed, and CFUs were enumerated at 0, 2, and 4 dpi.

### Bacterial strains

Bacterial strains were reported in (*17*), including *Lactobacillus plantarum WF*, *L. plantarum LF*, and *L. plantarum WCFS1*, which was called *L. plantarum HS* in (*17*). *Acetobacter indonesiensis* SB003 was assayed for colonization in Fig. S1 of (*17*). Fluorescent protein-expressing plasmid strains were developed and reported in (*17*) and (*44*). pCD256-p11-mCherry, used for *L. plantarum*, was the generous gift of Reingard Grabherr (BOKU, Austria) (*46*). pCM62, used for *Acetobacter indonesiensis*, was the generous gift of Elizabeth Skovran (SJSU, USA).

### Colonization assay

The colonization assay followed the protocol used in Fig. S1A of (*17*). Briefly, a measured dose of bacteria was pipetted evenly on the surface of a germ-free fly food vial and allowed to absorb for 15 min. 25 germ-free, 5- to 7-d post-eclosion, mated female flies were introduced to the vial and allowed to feed for a defined period of time. Flies were then removed from the inoculation vial and placed in fresh, germ-free vials. Bacteria were collected from the inoculation vial by vigorous rinsing with PBS, and the abundance was quantified by CFUs. At specified time points, CFUs in individual flies were enumerated by washing the flies 6 times in 70% ethanol, followed by rinsing in ddH_2_O, and then crushing and plating for CFU enumeration.

### Preparation of bacteria

Cultures of bacteria were grown overnight in 3 mL liquid media at 30 °C. *Lp* strains were grown in MRS liquid media (Hardy Diagnostics, #445054), and 10 µg/mL chloramphenicol was added for mCherry-expressing strains. *Ai* was grown in MYPL media, and 25 µg/mL tetracycline was added for GFP-expressing strains. Bacteria were pelleted by spinning for 3 min at 3000 rpm, resuspended in PBS, then diluted to the desired concentration. Dose size was quantified using OD_600_ or by plating and counting CFUs. OD of 1.0 corresponds to 2×10^8^ CFUs/mL for *LpWF* and 3×10^8^ CFUs/mL for *Ai*.

### Inoculation of flies

Flies were inoculated by pipetting 50 µL of an appropriate concentration of the inoculum onto the food and then left to dry in the biosafety cabinet for 15 min. Flies were starved for 4 h before flipping them into the inoculation vials, where they were allowed to feed for 1 h, then flipped to fresh vials. The dose per fly was calculated as the amount of inoculum consumed divided by the number of flies in the vial. To verify that flies ate the bacteria placed on the food and measure the amount of ingested inoculum, uneaten bacteria were recovered from the vial after feeding and subtracted from the original dose. For experiments to standardize the dose of bacteria, the vial was an inverted 50-mL conical vial with solidified agar food in the cap. This vial allows for separation of food CFUs from CFUs on the walls of the vial. For other experiments, the vial was an autoclaved, polypropylene wide fly vial (FlyStuff).

### Quantification of CFUs in flies

Abundance in the gut was measured by homogenizing whole flies then plating to count CFUs. Flies were first anesthetized using CO_2_ and surface-sterilized by washing twice in 70% ethanol, then twice in PBS. Next, they were placed individually into wells of a 96-well plate along with 100 µL PBS and ∼50 µL of 0.5-µm glass beads (Biospec) and heat-sealed (Thermal Bond Heat Seal Foil, 4titude). The plate was shaken violently for 4 min at 2100 rpm on a bead beater (Biospec Mini-beadbeater-96, #1001) to homogenize the flies. We previously showed that the 0.5-µm bead size does not diminish bacterial counts and effectively disrupts fly tissue (*17*). A dilution series of the entire plate was prepared using a liquid-handling robot (Benchsmart). Agar growth medium was prepared in rectangular tray plates, which were warmed and dried ∼30 min prior to plating. Plates were inoculated with 2 µL of fly homogenate per well, which leads to a circular patch for CFU enumeration. The plates were incubated at 30 °C overnight. To count colonies, plates were photographed under fluorescent light and counted semi-automatically using ImageJ.

### Measurement of CFUs in fly vial

The number of bacteria in a fly vial was measured by recovering cells from the vial and plating on nutrient agar growth media (MRS or MYPL) to count CFUs. To collect bacteria, 2 mL of sterile PBS were pipetted into the vial. The vial was then replugged and vortexed for 10 s. A dilution series was made starting with 100 µL of the PBS wash and then plated to count CFUs. This method was used to quantify viable bacteria egested (pooped) by flies, or bacterial growth in the vial or the remainder of uneaten inoculum. Egestion and inoculation were measured over a period of 1-2 h, minimizing the opportunity for new bacterial growth.

### CAFÉ assay

Twelve flies were placed in a sterile polypropylene wide mouth fly vial containing 2 mL of 1.2% agar in ddH_2_O. Four glass capillary tubes were inserted through the flug and filled with 12 µL of filter-sterilized liquid fly food (10% glucose, 5% yeast extract, 0.42% propionic acid). Ten microliters of overlay oil were added on top to push the liquid food to the bottom of the capillary. Flies were left in the vial for 24 h before being transferred to a fresh setup. Vials were checked every 12 h to ensure flies had access to food, and a fresh flug with new capillaries was inserted if capillaries had air in them, which prevents food access. Five fly vials were put together into a 1-L beaker with a wet paper towel at the bottom and aluminum foil over the top, and the beaker was placed in the back of a fly incubator set to 25 °C, 12 h-12 h light-dark cycling, and 60% relative humidity.

### Pulse-chase protocol for bacterial colonization

To estimate the turnover time of established bacterial populations, 5- to 7-day old mated female flies were kept with 25 flies/vial. Flies were first inoculated with a pulse of fluorescently labeled, antibiotic-resistant bacteria by pipetting 50 µL of culture resuspended in PBS (OD_600_=1) onto the food and allowing it to dry prior to flipping flies into the inoculation vial. The pulse dose was allowed to establish colonization in the gut for 3 d prior to chase. Flies were fed a chase dose in the same way each day for 10 d (OD_600_=1). The abundance of labeled resident was measured daily by homogenizing and plating a sample of flies on selective media to count CFUs. The invading chase dose was assayed by plating on non-selective media. To control for any other factors that might affect resident abundance, a control group was also passaged daily to fresh food with no chase dose and assayed daily to count CFUs.

### Pulse-chase analysis

Experiments were conducted in triplicate. Measurements from individual flies from the different experiments were pooled by timepoint. Data were fit to an exponential decay using prism, and the half-life with its confidence interval was reported.

### Measurement of growth rates *in vivo*

Plasmid loss in the absence of selection was used as a proxy for bacterial growth rate. Briefly, a standard curve was constructed by passaging plasmid-containing cells to fresh media twice daily in a ∼1:100 dilution to an OD of 0.01 for 6 d. The number of bacterial generations was estimated by counting the number of CFUs in the culture prior to dilution. The ratio of plasmid-containing CFUs to plasmid-free CFUs was counted as the number of fluorescent to non-fluorescent colonies. We note that the doubling time is roughly 2 h for each strain. A linear regression was used to fit an equation to the standard curve data. Flies were then fed 100% plasmid-containing cells. The ratio of plasmid-containing to plasmid-free CFUs was counted at various time points in the experiments, and the standard curve was used to convert the ratio to the number of doublings. In the case of dual-plasmid containing strains (Fig. S7C), growth was measured as a ratio of colonies positive for GFP-Erm plasmids (which are lost rapidly) to those positive for mCherry-Cam (which is retained much longer). A non-linear (exponential decay) regression was used. Two caveats we note are that (1) population bottlenecks cause wider variance in the plasmid ratio, and (2) *in vivo* plasmid loss rates may be different from *in vitro* rates. We previously showed that the first caveat, high variance due to bottlenecks, can be used to infer bottlenecks. We also note that with respect to the second caveat, our use of this method to compare growth rates in a controlled experiment does not necessitate an absolute growth measurement with a standard curve. Furthermore, the growth rates *in vivo* were similar to *in vitro*, meaning that any differences in plasmid loss rates due to differences in the growth phase of the cells are likely small.

### Cropectomy

Cropectomy was performed on live flies using only new, undamaged fine forceps (#5, Dumont). Forceps, flypad, and microscope area were cleaned with 70% ethanol. Five- to 10-day old female flies were first anesthetized using CO_2_ then placed on a depression slide for surgery. The fly was positioned on its back, and while holding the torso with one set of forceps a small puncture was made in the abdomen just below the thorax as shown in Fig. 1O. Pressure on the forceps was released slightly to allow the tips to open up, then grab onto the crop and pull it out through the puncture. If the crop duct was still attached, it was severed along the edge of a forceps. Flies were placed in a sterile food vial and given at least 3 d to recover. Survival rate was ∼1 in 10 flies.

### Preparation of samples for microscopy

Whole guts were removed from the fly by dissection with fine forceps (Dumont). Tissue was fixed in 4% PFA in PBS for 3 h at 24 °C or at 4 °C overnight. Guts were permeabilized using 0.1% Triton-X in PBS for 30 min at room temperature, washed twice in PBS, stained with 10 µg/mL DAPI for 30 min, washed twice in PBS, placed in mounting medium for up to 1 h, then transferred to the slide using a wide bore 200-µL pipette. Each gut was then positioned on a positively charged glass microscope slide, and approximately 60 µL of mounting medium was added (mounting medium: 80% glycerol, 20% 0.1M Tris 9.0, 0.4g/L N-propyl gallate). Five to ten 0.1-mm glass beads (Biospec) were added to the mounting medium to form a spacer that prevents crushing of the sample. The slide was then covered with a No. 1.5 cover glass and sealed with nail polish.

### Confocal microscopy

Microscopy was conducted with a Leica DMi8 confocal microscope using either a 40X (1.30 NA) HC Plan Apo or a 60X (1.40 NA) HC Plan Apo oil immersion objective. Laser lines were generated using a white-light laser with AOTF crystal, and excitation wavelengths for fluorophores were: mGFP5, 488 nm; mCherry, 591 nm; Cy5, 650 nm. Whole gut images were generated by tiling multiple captures then merging using the Mosaic Merge function in LAS X to stitch into a single stack. *Z*-stacks for whole guts were 70-80 µm in thickness with slices every 0.5 µm or less. To render two-dimensional images for publication, fluorescence channels were processed as maximum intensity *z*-projections and the brightfield channel is represented by a single *z*-slice from the middle of the stack.

### Measurement of fluorescence intensities

Fluorescence intensity of gut colonization was quantified using FIJI. Summed intensity *z*-projections of 80-µm optical sections were generated, then resized to a scale of 1 µm/px. Background subtraction with a rolling ball radius of 50 px was applied. A segmented line with spline fit and a width of 50 µm was drawn along the length of the gut, starting with the most distal point on the crop as the origin. The “Plot Profile” function was used to measure the intensity along each of 5 segments: crop, crop duct, proventriculus, midgut, and hindgut. Segment length was normalized to a standard length. Intensity was normalized by averaging each replicate then normalizing the means.

### Measurement of beads egested

To measure shedding of polystyrene beads (Spherotech FP-0552-2, sky blue), flow cytometry was used to quantify the number of egested beads. Flies were kept in inverted 50-mL conical tubes with 1 mL solid food in the cap. To collect shed material, the tubes were rinsed with 10 mL of PBS, vortexed for 10 s, and then a clean cap was placed on top. To concentrate the solution, the samples were spun in a centrifuge for 7 min at 3000 rpm. The pellet was then resuspended in 200 µL of PBS. The concentrated sample was counted on an Attune flow cytometer (Thermo Fisher).

### Electron microscopy

Whole guts were dissected in Cacodylate pH 7.4 (Cac) buffer, then fixed for 2 d in 3% GA+1%FA in 0.1 M Cac at 4 °C. Samples were embedded in agarose and stored at 4 °C until further processing. Samples were then washed in Cac buffer, stained with 1% OsO_4_+1.25% KfeCN for 1 h, washed in water, treated with 0.05 M Maleate pH 6.5 (Mal), stained with 0.5% Uranyl Acetate in Mal for 1.25 h, then washed with increasing concentrations of ethanol. For embedding in resin, samples were treated with resin+propylene oxide (1:1) evaporated overnight as a transition solvent prior to embedding, then embedded in epoxy resin (Epon+Quetol (2:1)+Spurr (3:1)+2% BDMA overnight at 55 °C and cured at 70 °C for 4 d.

### X-ray microcomputed tomography (XR µCT)

Samples were prepared for XR µCT following the protocol of Schoborg et al 2019 (*36*), which the authors generously shared prior to publication. Briefly, flies were washed in 1% Triton-X in PBS to reduce cuticular wax. A shallow hole was poked in the abdomen and thorax with a fine tungsten pin to increase permeation of fixative and stain. Fixation was with Bouin’s solution. Staining was with phoshotungstic acid for 3 weeks. Flies were mounted for imaging in a 10-µL micropipette tip containing deionized water and sealed with parafilm. Imaging was performed at the Lawrence Berkeley National Laboratory’s synchrotron Advanced Light Source on beamline 8.3.2 with assistance of Dula Parkinson. 1313 images were acquired per specimen at 20X magnification through 180 degrees of rotation. Back-projections were performed using Tomopy with the following specifications:

*doFWringremoval 0 doPhaseRetrieval 1 alphaReg 0.5 doPolarRing 1 Rmaxwidth 30 Rtmax 300*

Further specifications are available here: http://microct.lbl.gov/. The images in Figures 6A,B were produced in Octopus. Volumetric reconstructions of the gut lumen in Figure D-G were performed in Imaris using manual segmentation.

### Statistics

Statistical tests were performed in Prism. In general, data were checked for normality using a Shapiro-Wilk test. If normality was established, a Welch’s t-test was performed. Statistical tests of CFU abundances were performed on log_10_-transformed data. When CFUs were 0, the log was set to 0 (corresponding to a pseudocount of 1). When multiple comparisons were made, an ordinary one-way ANOVA was performed. If significant, multiple pairwise comparisons were performed with Tukey’s multiple comparisons test. When data was not normally distributed, comparisons were made using Wilcoxon rank-order tests. Error bars on proportions are either standard error of the proportion (s.e.p.), or binomial 95% confidence intervals using the Clopper-Pearson method or Jeffries method, as specified in the text. The statistical significance of differences in proportions were assessed using a Z-test.

## Supplementary Text

### *Ai* exhibits increased early death rates in germ-free flies

To probe the facilitation of *Ai* colonization by *LpWF*, we examined the dynamics of *Ai* colonization from 1 hpi to 6 dpi (Fig. S2F, S5F-M, S8). For the first 1 dpi, *Ai* abundance was significantly higher in *LpWF*-pre-colonized versus germ-free flies (Fig. 3A, S5F). After 2 dpi, *Ai* levels were only slightly higher in *LpWF*-pre-colonized flies (Fig. S5F). Thus, the presence of *LpWF* ameliorates the initial decrease in *Ai* levels, which could stem from a decrease in the growth rate, an increase in the death rate or the egestion rate, or some combination of these factors. We comprehensively measured each of these rates.

We measured growth rate in the fly using fluorescent protein plasmid dilution due to growth in the absence of antibiotic selection (Fig. S5B-H) (*17*). The mean generation time of *Ai* was similar in initially germ-free and *LpWF*-pre-colonized flies (0.21 vs. 0.23 h^-1^, Welch’s t-test, *p*= 0.75; Fig. 5B). However, the variance in plasmid loss was significantly higher in germ-free flies compared with *LpWF*-pre-colonized flies (F-test, *p*=0.014), consistent with the observed population bottleneck (Fig. S5F), which we also previously observed in certain *Lp* strains and connected to a population bottleneck shortly after inoculation (*17*). Thus, different growth rates of *Ai* cells with or without *LpWF* do not seem to account for the differences in *Ai* abundance.

To determine whether the initially germ-free flies egested *Ai* cells more rapidly than *LpWF*-pre-colonized flies, we measured the egestion rate from the abundance of *Ai* in their frass (excrement) after 1 h in a fresh vial. The rate of viable *Ai* egested by initially germ-free flies reached zero by 1 dpi, while *Ai* egestion in *LpWF*-pre-colonized flies remained higher and never reached zero (Fig. S8). Differences in egestion rate could be due to more rapid passage through the fly or to variable death rates of the bacteria inside the fly. To measure rates of passage through the fly, we fed fluorescent polystyrene beads simultaneously with *Ai* inoculation, and the proportion of egested beads was quantified over time by flow cytometry (Fig. S5J). The rate of bead egestion was highly similar between *LpWF-*pre-colonized and germ-free flies (Fig. S5J). Thus, transit time through the gut does not explain the differences in *Ai* colonization dynamics, suggesting a higher death rate of the *Ai* cells colonizing an initially germ-free gut.

Since egestion is tightly linked to ingestion (*43*), we measured the total *Ai* consumed by flies versus that remaining in the vial after feeding by counting CFUs in flies and on the food 1 hpi, reasoning that any bacteria not accounted for must have died during the 1 h of feeding (Fig. S5, S9), e.g. by lysis in the digestive tract. In both sets of flies, only a small fraction of the inoculum was left 1 hpi (Fig. S1 , S9). These measurements indicate that germ-free and *LpWF*-pre-colonized flies consumed the same amount of *Ai* and that *Ai* has a higher survival rate in the gut of *LpWF*-colonized flies.

The higher survival in co-colonized guts could be due to bacterial interspecies interactions, such as a cytoprotective effect of *LpWF* on *Ai*, or to host-microbe interactions, such as the fly gut becoming more hospitable to *Ai* when pre-colonized by *LpWF*. To differentiate between these two possibilities, we fed germ-free flies with *LpWF* and *Ai* simultaneously, reasoning that host priming would not be evident with simultaneous colonization (Fig. S5L,M). *Ai* abundance at 1 hpi in co-inoculated flies was similar to initially germ-free flies fed *Ai* alone, and significantly lower than in *LpWF*-pre-colonized flies 1 hpi (Fig. S5M), indicating that *LpWF* remodels the host in a manner beneficial to *Ai*. We also measured *Ai* survival 1 hpi when colonizing *Ai*-pre-colonized flies. A slight advantage was observed (Fig. S5L), which was substantially less than for *Ai* colonizing *LpWF* flies (c.f. Fig. 3A). *In vitro*, *Ai* abundance was unaffected by co-culturing with *LpWF* (*44*). Because the *Ai* cells are alive in the proventriculus but dead upon defecation, a simple explanation consistent with our data is that for *Ai* cells colonizing *LpWF*-pre-colonized flies, more *Ai* cells are retained for a longer period of time in the proventriculus, and cells that are not retained in the proventriculus die when passing through the midgut. Taken together, our results indicate that the host environment is more permissive to *Ai* survival when pre-colonized by *LpWF*.

## Supplementary Figures (S1 through S9) follow

**Figure S1.**
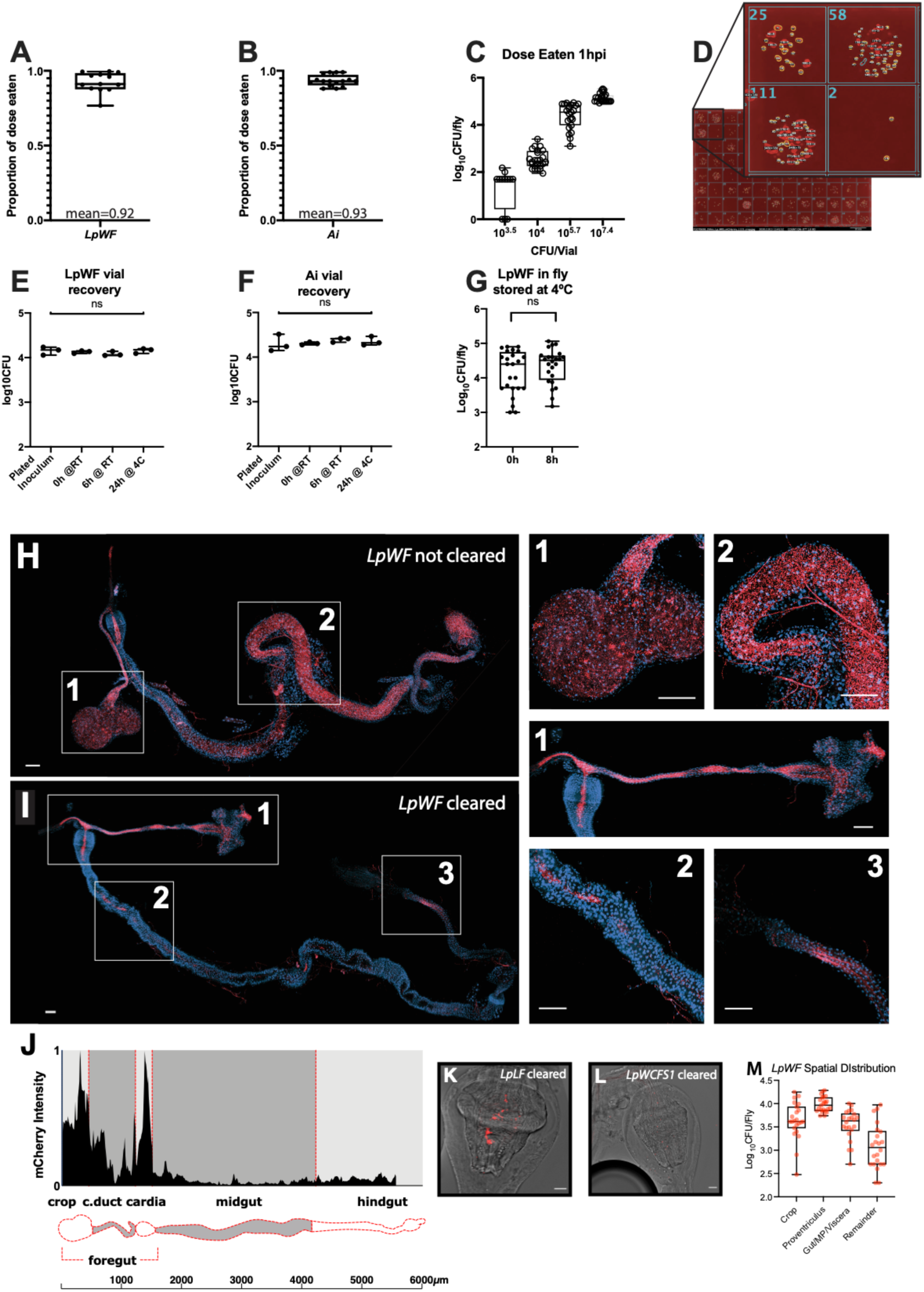
Validation of colonization assay and culturing techniques. A. *LpWF* dose consumed was assayed by washing the food in 1x PBS and plating the solution. A dose of 10^3.5^, 10^4^, or 10^5.7^, 10^7.4^ CFUs/vial was fed on top of agar food in standard vials. Flies ate >90% of the dose after 1 hour (n=12 vials, mean=0.9235). Results were normalized to the dose. The proportion of the dose consumed was calculated by subtracting the leftover inoculum from the delivered inoculum and normalizing to the delivered inoculum. The growth of bacteria on the food over the 1 hour feeding window was monitored by using a parallel control vial that did not have flies added (see panel E). B. *Ai* dose eaten: flies ate >90% of dose after 1 hours (n=16 vials, mean=0.93). Same methods as panel A. C. CFU abundance in flies 1 hpi. Flies were inoculated by feeding on standard food, 25 flies/vial. For doses 10^3.5^ , 10^4^, 10^5.7^, and 10^7.4^ CFUs/vial (equal to 10^2.1^ , 10^2.6^ , 10^4.2^ , and 10^6^ CFUs/fly respectively), flies all ate a similar amount of bacteria. For the lowest dose, 10^3.5^ CFUs total in the vial, which was about 125 CFU/fly, 3 of 12 flies sampled had 0 detectable CFUs 1 hpi. The limit of detection was 50 CFUs. D. For CFU quantification, flies were collected into 96 well plates containing 100µl PBS and 0.1µm glass beads. In our standard assays, CFUs were quantified by spotting 2µl of the 100 µL fly homogenate (in 96 well plates) onto growth media in rectangular tray plates so that each well of the 96 well plate was spotted. Microcolonies were grown for 30 h at 30°C. Counting was performed by photographing plates, counting colonies in ImageJ, and manually validating. Because the maximum amount of homogenate plated is 1/50th of a fly, a count of 1 colony yields a value of 50 CFUs/fly; the *resolution* of this quantification system is 50 CFUs, which we also call the limit of detection (LOD). To distinguish the invading strain from the resident strain in the priority effects experiments, invading bacteria containing a resistance plasmid were used and plated on selective media, CFU quantification in GF control flies was done in parallel during the same experiment using also the same plasmid-containing inoculum and counted on the same selective media. E. Validation experiment shows that the number of CFUs recovered did not vary significantly from the inoculum measured by directly plating. Bacteria were recovered from vials by rinsing with 2mL PBS then plating a dilution to count CFUs. Inoculum was recovered immediately after placing on the fly food, after leaving at room temperature for 6 hours, and storing at 4°C overnight. *LpWF* bacteria were used. F. Validation of *Ai* recovery from vials was the same as in D, CFU counts were consistent for *Ai*. G. When flies could not be homogenized and plated immediately, they were stored at 4°C for up to 8 h. To test for any possible effects on the bacterial abundance, flies from the same vial were homogenized either immediately or after storage for 8 h at 4°C (n=23 flies/time point). There was no significant difference in CFU counts. (n=46, unpaired t-test p=0.2794) H. Transient microbes are found throughout the gut in flies kept on the same food for more than 24 h. *LpWF* is labeled with mCherry (red). Note the distended crop (1) and food filled midgut (2), the punctate appearance of the mCherry indicates bacteria dispersed throughout the fly food. Blue is a single z-slice of DAPI stain to indicate the gut boundary. I. Guts were cleared of transient microbes by placing on agar-water starvation media for 3 hours. *LpWF* remains in the foregut (1), the esophageal tract is lined with a dense and continuous population of *LpWF*, whereas there is a patchy appearance in the crop. mCherry signal is largely absent from the midgut aside from a few small patches (2), and it is absent from the hindgut, although some autofluorescence occurs (3). J. Quantification of spatial distribution of *LpWF* in the fly digestive tract. Mean intensity of mCherry fluorescent signal in *LpWF*-mCherry-colonized flies 3-5 dpi, n=5 guts. Drawing depicts a segmentation of an average gut oriented lengthwise beginning with the crop. Intensity along length was normalized to the standard length per region. K. Microscopy of *LpLF* in the proventriculus. Scale bar 20 µm. L. *LpWCFS1* in the proventriculus. Scale bar 20 µm. M. Raw CFU counts of spatial distribution of *LpWF* in dissected gut regions (Fig. 1G) shows the majority of CFUs in the fly gut are in the proventriculus and crop duct.

**Figure S2.**
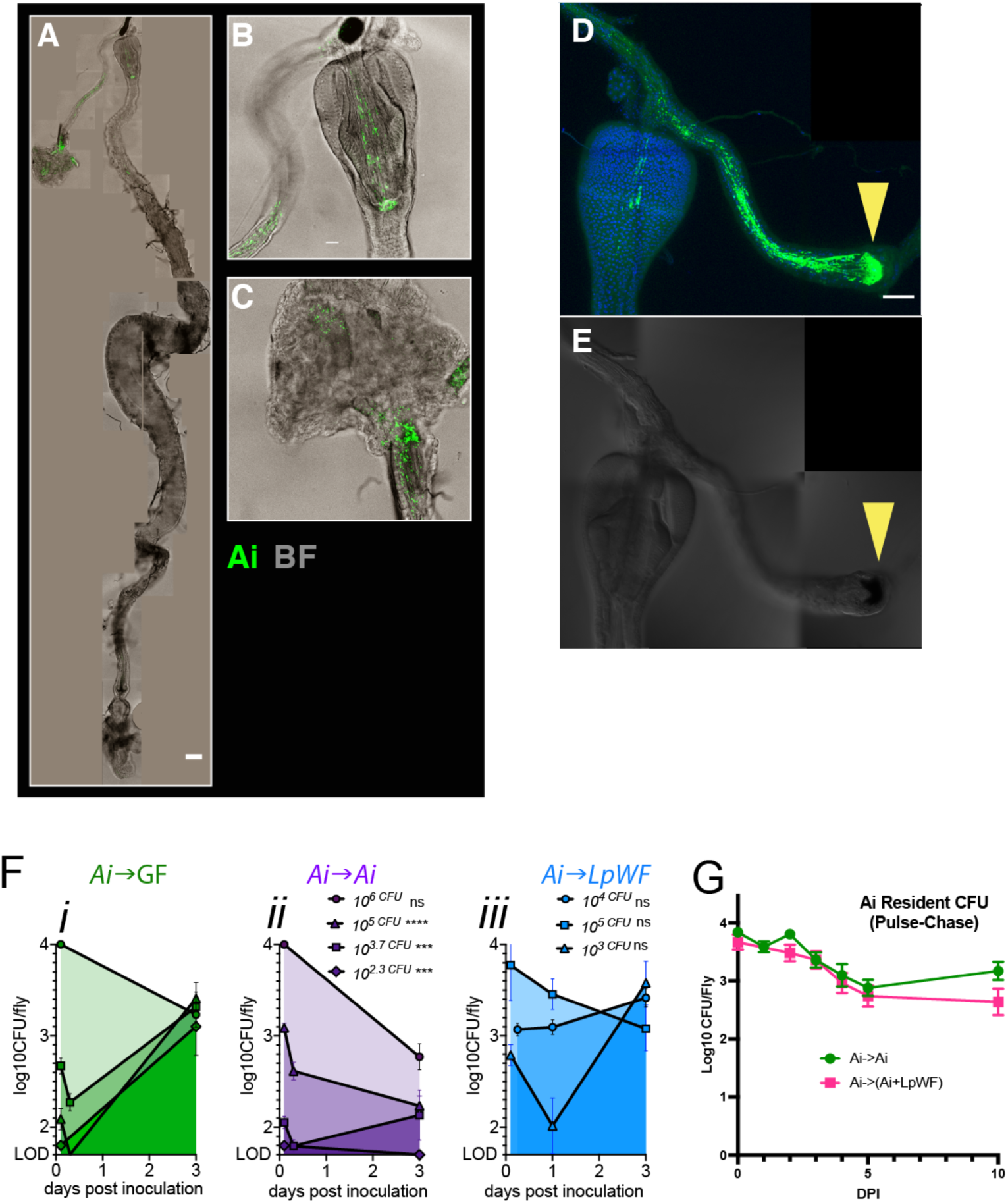
*Ai* colonization is similar to *LpWF* colonization. A. Whole mount gut colonized by *Ai-*mGFP5. B. Detail of proventriculus. C. Detail of crop. D. *Ai* colonization of the foregut after cropectomy surgery. Green = *Ai*-mGFP5. Blue = DAPI. Scale bar 50 µm. n=14 of 14 flies colonized after cropectomy surgery. E. Brightfield image of the foregut in D. Yellow arrowhead indicates melanization at site of crop duct severing. F. Time course bacterial abundance for *Ai* colonizing (i) germ-free, (ii) *Ai*-colonized, and (iii) *LpWF*-colonized flies. n=24 flies per time point. G. Pulse-chase of *Ai* into *Ai-*mGFP5-pre-colonized flies (green) or flies pre-colonized by *Ai-*mGFP5 and *LpWF* (pink).

**Figure S3.**
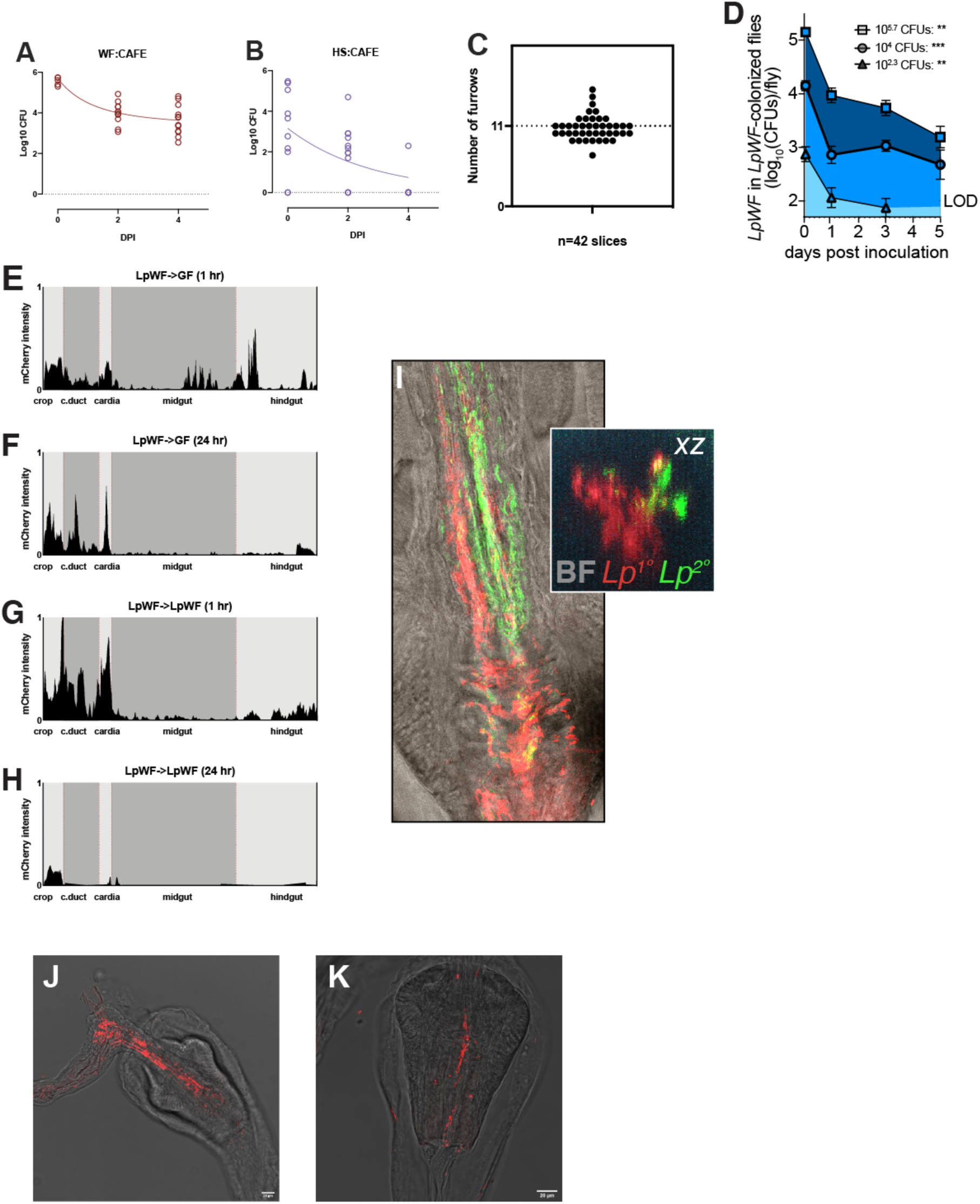
Timecourses of microbe population abundance in the gut. <u>Doses:</u> *LpWF* Low: 2.0 x 10^3^ CFUs/fly; *LpWF* High: 5.7 x 10^5^ CFUs/fly. 12 flies were sampled daily and analyzed for CFU counts. A. *LpWF* High into GF flies transferred daily to a fresh vial with only CAFÉ-supplied liquid food (10% glucose, 5% yeast extract, 0.42% propionic acid) over 5 dpi. B. *LpWCFS1* High into GF flies transferred daily to fresh vial with only CAFÉ-supplied liquid food (10% glucose, 5% yeast extract, 0.42% propionic acid) over 5 dpi. C. The number of furrows in the proventriculus, mean=10.89. Furrows were counted in 42 TEM images from various points along the length of the proventriculus. n=5 different proventriculi. D. A single dose of *LpWF-*mCherry was fed at a range of doses (see inset) to flies pre-colonized by *LpWF* and *LpWF*-mCherry CFUs were quantified over 5 d, indicating the abundance does not converge at ∼10^4^ CFUs/fly as when the doses are fed to initially germ-free flies (Fig. 2A). E. (E-H) Quantification of spatial distribution of *LpWF* along the gut. Mean intensity of *LpWF*-mCherry fluorescence fed to either GF or *LpWF* pre-colonized flies at 1 hour or 24 hours after inoculation. Summed intensity projections of 80-µm thick stacks of confocal images of whole gut dissections were quantified for fluorescence intensity, normalized to total intensity and length. N=5 flies per treatment. E: *LpWF*-mCherry → GF at 1 hpi. F. *LpWF*-mCherry → GF at 24 hpi. G. *LpWF*-mCherry → *LpWF* at 1 hpi. H. *LpWF*-mCherry → *LpWF* at 24 hpi. I. *LpWF-*sfGFP → *LpWF-*mCherry 1 hpi of *LpWF-*sfGFP. Confocal fluorescent image of proventriculus. Inset: optical x-z cross section. Note that we rarely observed the secondary colonizer in the furrows, but when we did, the cells of the secondary dose clustered tightly.

**Figure S4.**
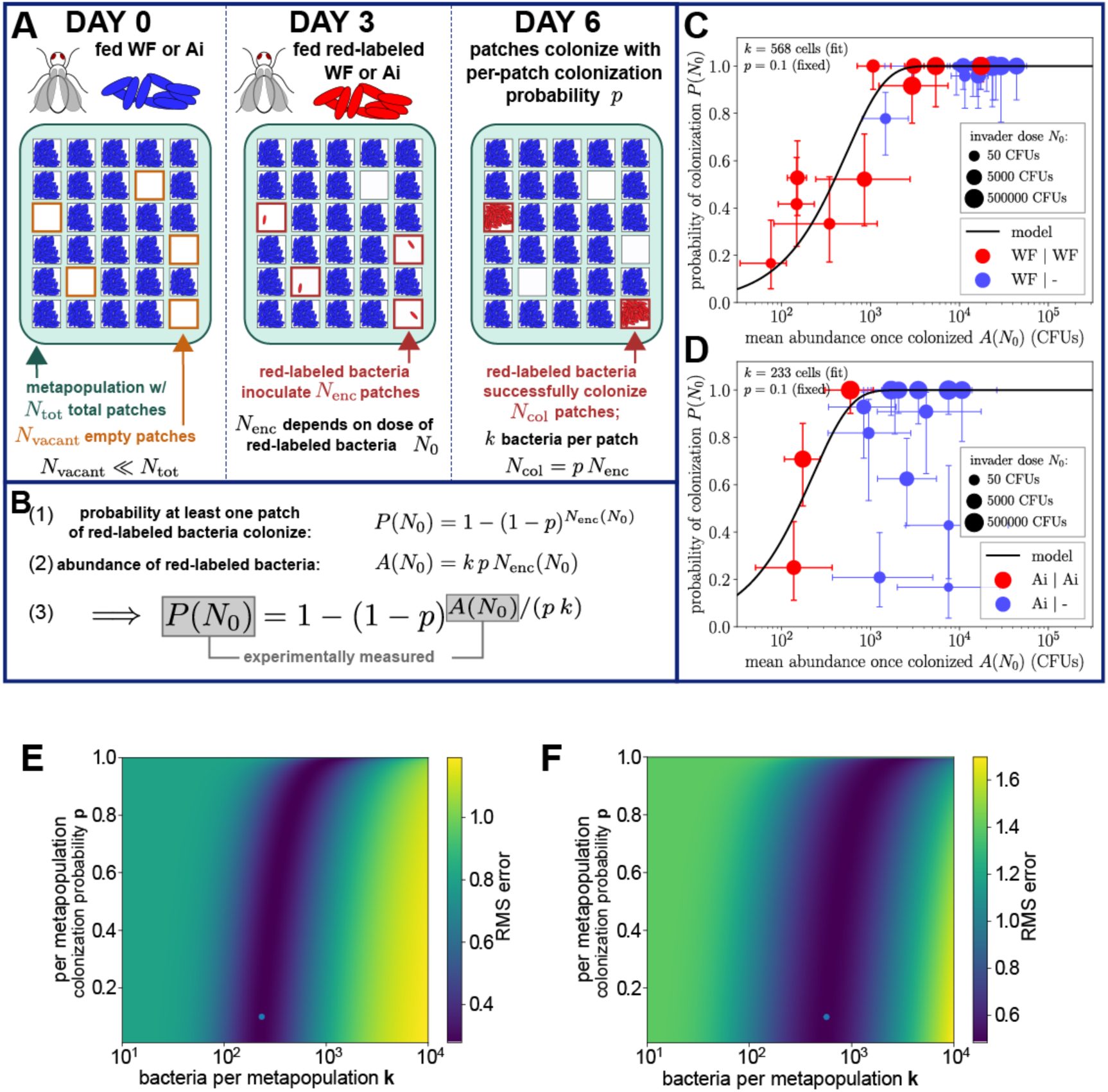
Quantitative model of colonization relates final colonized abundance of invader to the probability of colonization at the single fly level. A. Model describes the spatial variability of bacterial colonization with a metapopulation model of patchy colonization, assuming that the fly gut may be subdivided into 𝑁*_tot_* subpopulations based on the observation that turnover occurs on the time scale of 15 d. On day 0 a strong colonizer (*LpWF* or *Ai*, colored blue) is fed to the fly. By day 3, the initially fed blue bacterial species are assumed to colonize the majority of the patches, leaving 𝑁*_vacant_* patches uncolonized. When colonized, each patch has a carrying capacity of k bacteria. On day 3 a red-labeled but otherwise identical bacteria (*LpWF* or *Ai*, colored red) is fed to the fly at an abundance 𝑁_0_, and the red-labeled bacteria proceed to inoculate some 𝑁_enc_ of these patches; with probability 𝑝 these inoculated patches become fully colonized with 𝑘 bacteria, and with probability 1 − 𝑝 they go extinct by day 6. B. Equation describing the model. (1) The probability of invader colonization as a function of the dose. (2) Abundance (𝐴) of invader in terms of the per-patch carrying capacity 𝑘, the per-patch probability of colonization 𝑝, and the number of inoculated patches 𝑁_enc_. Eliminating 𝑁_enc_ yields the third equation. (3) Relationship between the experimentally measurable probability of colonization 𝑃*_col_* and the invader abundance 𝐴. The two free parameters 𝑝 and 𝑘 may be fit; these parameters have the biological significance of indicating how bacteria are distributed among patches when colonizing, thus informing their spatial distribution. C. Consistent with the model, *LpWF* → *LpWF* priority effect experiments show a positive correlation between mean abundance 𝐴(𝑁_0_) and probability of colonization 𝑃(𝑁_0_), and when fit to the metapopulation model with 𝑝 = 0.1 fixed predicts the per-patch carrying capacity 𝑘 to be 568 cells D. *Ai* → *Ai* priority effect experiments predict a per-patch carrying capacity of 233 cells. Error bars show 95% confidence intervals (errors in probability of colonization computed with the Jeffreys interval; errors in mean abundance computed by bootstrapping log-transformed abundances). E. Error probability function for the fit of 𝑘 to the *LpWF* data shows that the fit of 𝑘 is robust. F. Error probability function for the fit of 𝑘 to the *Ai* data shows that the fit of 𝑘 is robust.

**Figure S5.**
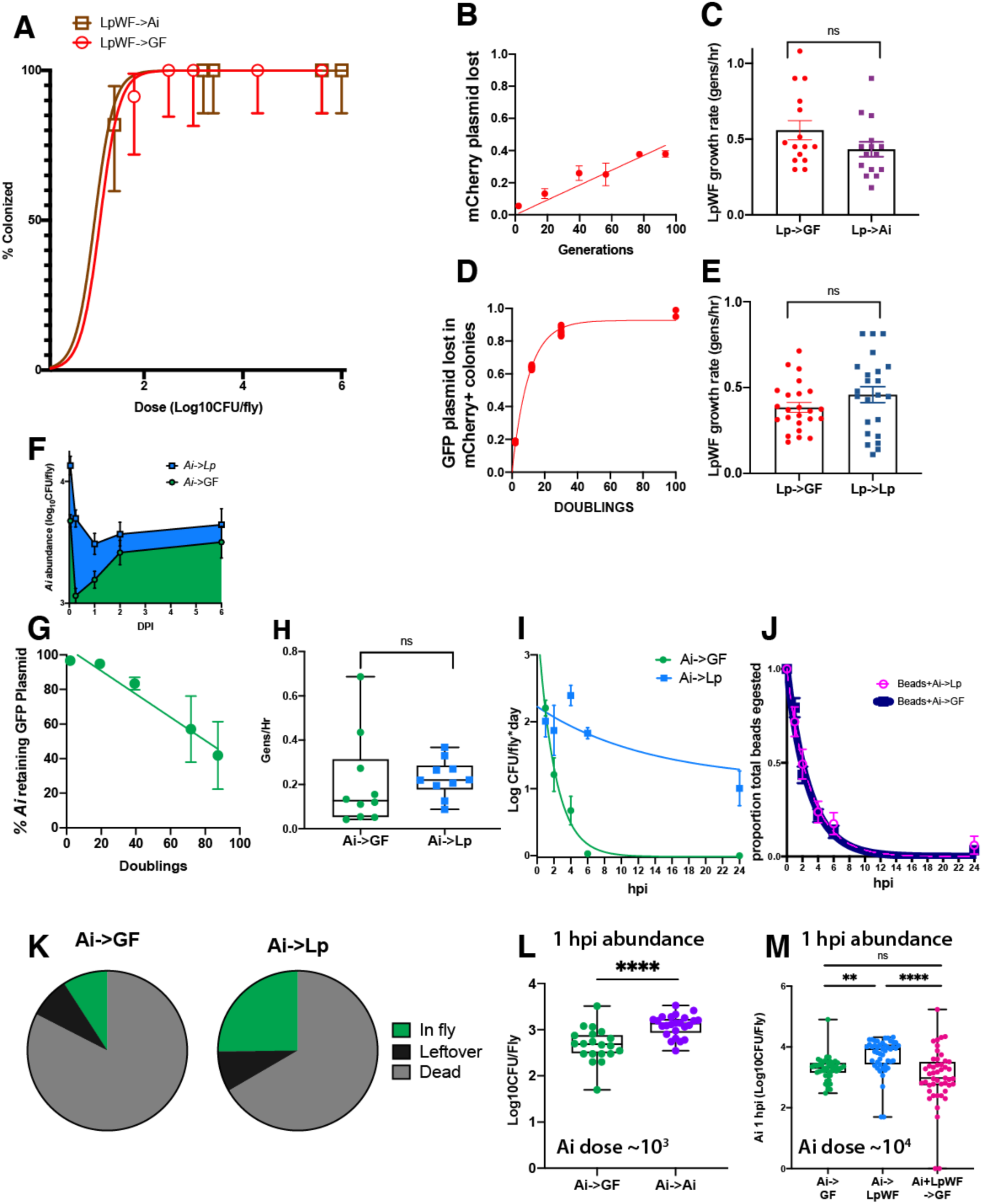
Dose response and kinetics in vivo. A. Dose response for *LpWF* fed to *Ai*-pre-colonized flies. B. Plasmid loss standard curve for pCD256NS-P1-mCherry-ΔEc (mCherry-Cam) in *LpWF* enumerated daily for 5 d by plating on non-selective media and counting fluorescent vs. non-fluorescent colonies. 100-fold daily dilution in 3 mL culture. Slope of the simple linear regression of *in vitro* plasmid loss rate was 0.004624 of total colonies per doubling event (R^2^=0.07301). C. Growth rate of *LpWF in vivo*. *LpWF* invading GF flies had a mean growth rate 0.4589 gen/hr 5 d after invasion while *LpWF* invading *Ai* flies grew at a lower but not significantly different mean of 0.3849 gen/hr (n=12 individual fly homogenates per condition; Welch’s t-test, p=.1272; F-test no significant difference in variances p=0.3499). The ∼2-fold variation in individual fly measurements is expected due to a population bottleneck that we previously chracterized (*17*). D. Dual plasmid standard curve: plasmid loss in *LpWF* containing both plasmids pCD256NS-P11-mCherry-ΔEc and pTRKH2-mGFP5 (GFP-Erm) was measured as a ratio of colonies positive for GFP-Erm plasmids (which are lost rapidly) divided by those positive for mCherry-Cam (which is retained much longer). This standard was modeled as an exponential function with a plateau: y = 1-(0.9326*exp(−0.07325*x)), R^2^=0.9986. E. Growth rate of *LpWF* invading *LpWF* pre-colonized flies. *LpWFCam/Erm* invading GF flies had a mean growth rate 0.4589 gen/hr, whereas *LpWFCam/Erm* invading *LpWF* pre-colonized flies had a mean of 0.5596 gen/hr as estimated from CFUs in flies 5 dpi with *LpWFCam/Erm*. There was no significant difference in growth rates (Welch’s t-test, p=0.1768). An F-test to compare variances was significant (p=0.034) where *LpWF* invading *LpWF* pre-colonized flies had a higher variance in plasmid loss, suggesting a founder effect due to lower initial population. F. *Ai* CFU abundance over time comparing flies germ-free at 0 dpi with flies pre-colonized by *LpWF* at 0 dpi. *Ai* abundance is lower in GF flies vs in flies pre-colonized by *LpWF* at 1 hpi, 6 hpi, and 1 dpi (*p*<0.0001, independent, unpaired t-tests, Bonferroni correction) but not at 2 dpi or 5 dpi (p>0.05) G. *Ai* plasmid loss standard curve: Growth in the absence of antibiotic selection leads to plasmid loss that is correlated with the number of cell divisions. The ratio of colonies with:without plasmid pCM62-mGFP5-tet (GFP-Tet) in *Acetobacter indonesiensis* SB003 was quantified daily for 5 d by plating on non-selective media and counting fluorescent vs. non-fluorescent colonies as a function of the total amount of culture growth. The slope of the linear regression of this standard curve was 0.56% percent of cells lost their plasmid every doubling event. This rate was applied to plasmid loss by bacteria in flies to estimate the *in vivo* growth rate. Percentage of plasmid was measured daily for 5 d. Y = 0.005579*x, (R^2^=0.1590) H. Mean growth rate 6 d after inoculation was 0.2287 generations per hour (gen/hr) for *Ai* invading *LpWF* pre-colonized flies or 0.2060 gen/hr for Ai invading GF flies (n=10 samples of 8 flies each). There was no significant difference in growth rates between *Ai* growth rate in flies (Unpaired Welch’s t-test, p=0.7528). Higher variance was observed for *Ai* invading GF flies (F-test, p=0.014). I. Transit time of *Ai* through the gut to GF or *LpWF*-pre-colonized flies in the first day after inoculation. *Ai* was fed along with polystyrene beads to flies (dose = 1.2 x 10^5^ CFUs of *Ai*/fly) in standard food in the cap of a 50 mL Falcon tube. *Ai* shedding was measured by counting CFUs recovered from falcon tubes by rinsing with PBS then centrifuging the contents to concentrate bacteria and beads for flow cytometry. Half-life of *Ai* in GF during the first day was 1.5 hours, while egestion of *Ai* in *LpWF* never decayed to zero. J. Shedding of 0.5-µm fluorescent, polystyrene beads co-fed to flies with *Ai* in FIG 5C. Beads were counted by flow cytometry. (∼4 x 10^5^ beads fed per fly). Half-life of beads was 1.9 or 2.0 hours in GF flies vs. in *LpWF* pre-colonized flies respectively, a non-significant difference (95% CI of decay fit). K. Proportion of *Ai* dose remaining in vials after feeding, viable in flies, or killed, n=12 vials per condition. Proportions are normalized among 3 groups of flies fed doses of 3.0 x 10^3^, 3.0 x 10^4^, and 3.2 x 10^5^ CFU/fly. L. Number of live CFUs of *Ai* in flies 1 hpi comparing *Ai* infor GF flies vs *Ai* into flies pre-colonized by *Ai*. Dose was ∼10^4^ CFUs/fly. n=20 flies/condition. M. Number of live CFUs of *Ai* in flies 1 hpi comparing *Ai* alone into GF flies versus *Ai* alone into *LpWF*-pre-colonized flies versus *Ai*+*LpWF* mixed into GF flies. Dose was 3 x 10^4^ CFUs of *Ai*/fly (n=48 flies/condition). For *Ai*+*LpWF* mixed, dose of *LpWF* was 3 x 10^4^ CFUs/fly.

**Figure S6.**
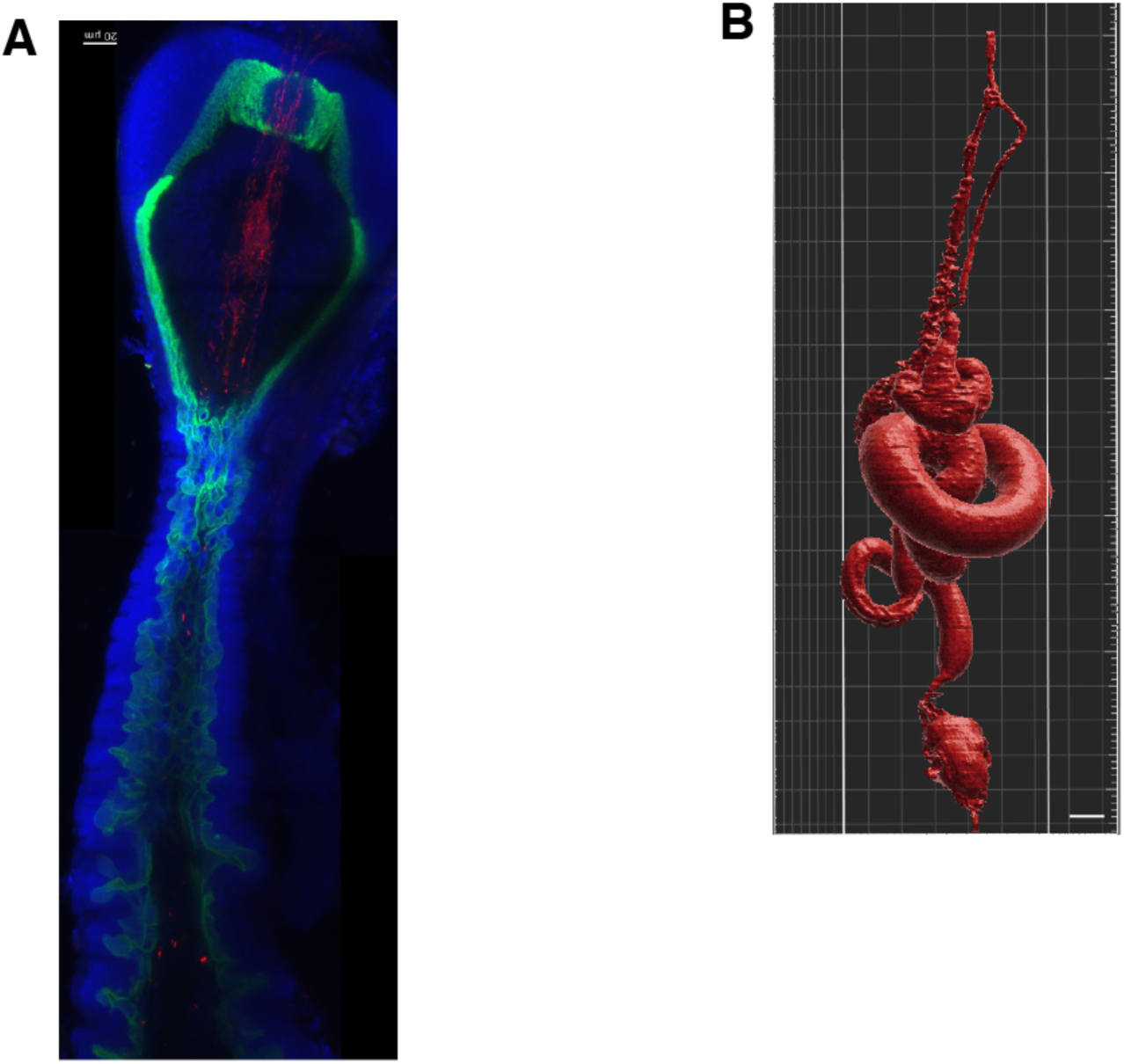
Imaging crypt spaces. A. Foregut colonized by LpWF-mCherry in A142-GFP brush border reporter transgene flies provided by the Buchon Lab. Brush) borders (green), *LpWF*-mCherry (red), DNA/DAPI (blue). Scale bar = 20µm. B. Whole fly gut model made using XR-µCT, as in FIG 6A-C. Used to compute volume of the 3 segments assayed in FIG 1N. Segment volume: Foregut: 5.08×10^6^ µm^3^, Midgut: 4.60×10^7^ µm^3^, Hindgut: 6.45×10^6^ µm^3^, Cardia: 3.39×10^5^ µm^3^, Crop: 4.75×10^6^ µm^3^, Visera: 5.24×10^7^ µm^3^. Rough surfaces in the volume rendering correspond to crypts that are visualized by the brush border marker in A.

**Figure S7.**
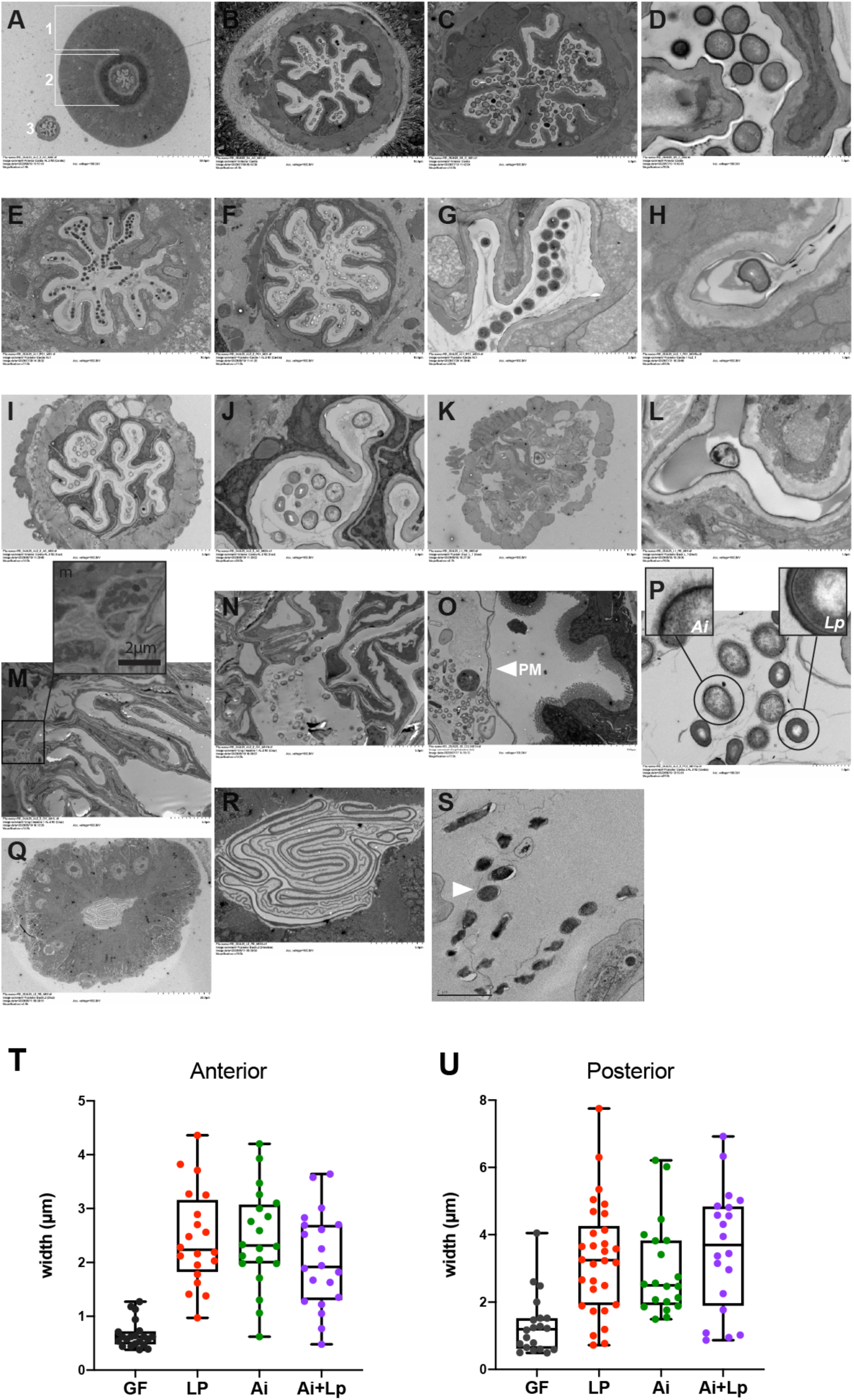
Spatial Structure of Colonization by Transmission Electron Microscopy. A. Overview of the anterior proventriculus. The mesodermal, midgut portion of the proventriculus (the proventriculus or outer proventriculus) is indicated (1). The ectodermal, foregut portion (the inner proventriculus or stomadeal valve) in indicated (2). The crop duct is present in this section as well (3). (*Ai+LpWF* colonized) B. Anterior proventriculus post feeding (*LpWF* 1 hpi) C. Anterior proventriculus post feeding (*LpWF* 1 hpi) D. *LpWF* packed in anterior proventriculus furrow (*LpWF* 1 hpi) E. Posterior proventriculus colonized (*Ai->LpWF* 1 hpi F. Posterior proventriculus colonized (*Ai+LpWF* 5 dpi) G. Posterior proventriculus furrow, (*Ai->LpWF* 1 hpi). Only *LpWF* visible. H. Long narrow furrow with single *Lp* cell (*Ai+LpWF* colonized) I. Crop Duct, similar morphology to proventriculus. (*Ai+LpWF* colonized) J. Detail of crop duct in I (*Ai+LpWF* colonized) K. Posterior crop duct/anterior crop, sparsely colonized (*LpWF* colonized) L. Single bacterium in posterior crop duct (*LpWF* colonized) M. Crop wall cuticle. Inset: cluster of bacteria. (*Ai+LpWF* Colonized) N. Crop lumen and cuticle (*Ai+LpWF* Colonized). O. Midgut, bacteria are separated from the brush borders (BB) by the peritrophic membrane (PM) (LpWF 1 hpi). P. Posterior proventriculus: both *Ai and LpWF* in the lumen of the posterior proventriculus. The gram negative *Ai* can be identified by a fuzzy coat (the glycocalyx or fimbriae) and its larger size relative to *LpWF. LpWF* is gram positive, it is distinguished by its think cell wall. (*Ai+LpWF* colonized) Q. Constriction between posterior proventriculus and anterior midgut, where the peritrophic matrix (PM) is extruded from proventriculus outer lumen (*LpWF* colonized). R. PM immediately posterior to the proventriculus (*LpWF* colonized). S. High pressure freezing shows cleared zone between the lumen wall and bacteria, indicating the boundary region shown in FIG 6P-2 is not a fixation artefact. T. Quantification of proventriculus furrow width in the anterior proventriculus for germ-free flies and flies colonized with *LpWF*, *Ai*, or *Ai+LpWF*. n=2 proventriculi per treatment and 10 furrow measurements per proventriculus. U. Quantification of proventriculus furrow width in the posterior proventriculus for germ-free flies and flies colonized with *LpWF*, *Ai*, or *Ai+LpWF*. n=2 proventriculi per treatment and 10 furrow measurements per proventriculus.

**Figure S8.**
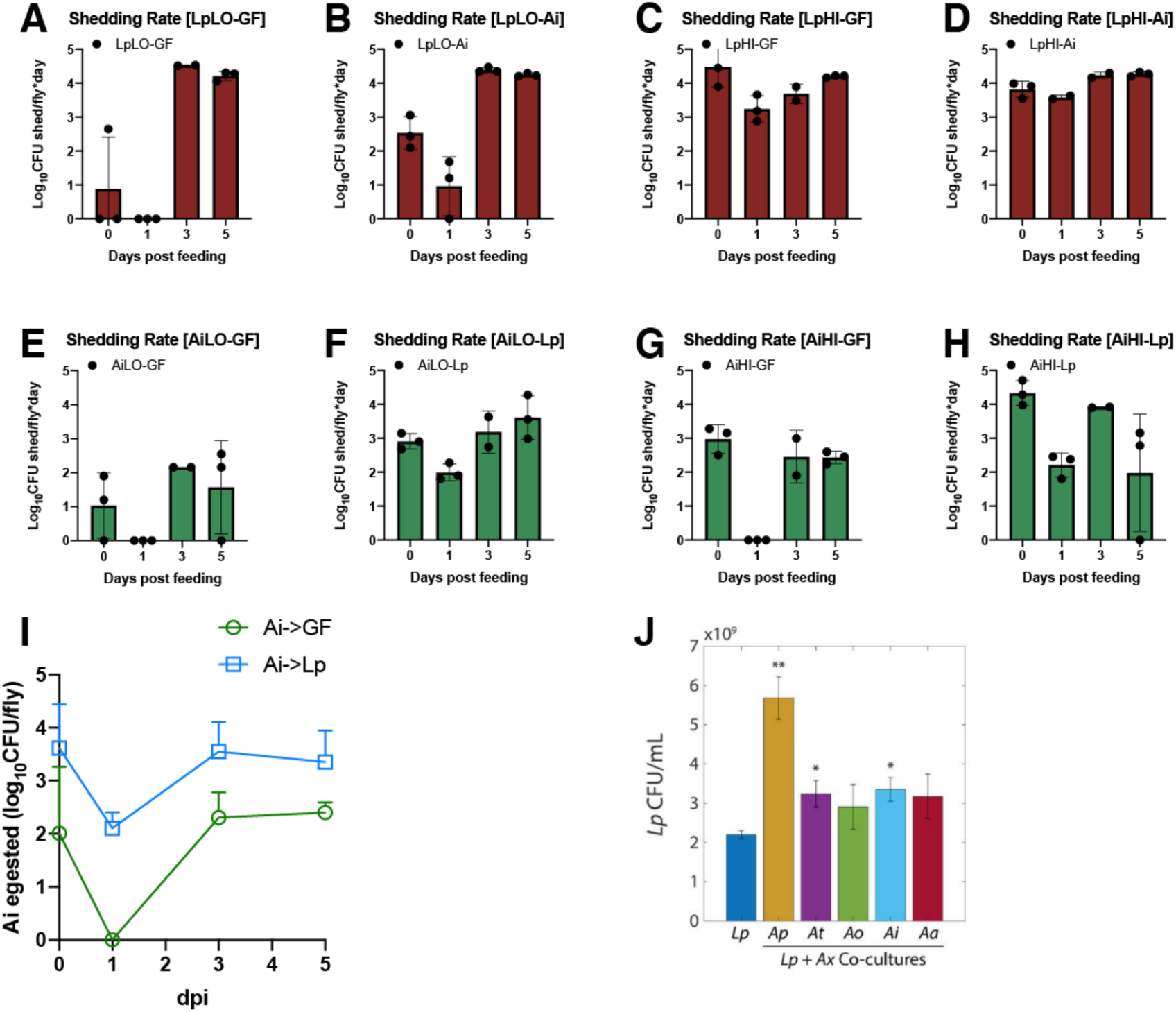
Egestion of bacteria by flies following inoculation. Shedding rates for various conditions following inoculation with bacteria were measured by keeping flies in a vial for a period of 1 hour, recovering viable bacteria from the vial by rinsing with PBS, then plating to count CFUs. Treatments correspond to the same experiments as in figures S5A-S5H. A. *LpWF* Low into GF flies. B. *LpWF* Low into flies pre-colonized by *Ai*. C. *LpWF* High into GF flies. D. *LpWF* High into flies pre-colonized by *Ai.* (A-D) Regardless of dose, *LpWF* egestion rate was lowest 1 dpi, suggesting a period of establishment. 3 dpi, *LpWF* CFUs are shed at a consistent rate of 2×10^4^ CFU/fly/day, about equal to the stable population of LpWF (FIG S5A). E. *Ai* Low into GF flies. F. *Ai* Low into flies pre-colonized by *LpWF* G. *Ai* High into GF flies. H. *Ai* High into flies pre-colonized by *LpWF*. (E-G) *Ai* shedding rate is variable over time and between treatments. I. Combined data from E-H plotted on same graph. After 24 hours, the average number of *Ai* egested reaches 0 in GF flies then increases to a mean of 2.5 x 10^2^ CFU/fly/hour The number of egested *Ai* in *LpWF*-pre-colonized flies is significantly higher at all time points, never drops to 0, and achieves an average rate of 3.2 x 10^3^ CFU/fly/day. J. Co-culturing *Lp* with *Ap*, *At*, *Ai*, or *Aa* resulted in increased *Lp* cell density after 48 h. Co-culturing with Ao did not significantly increase *Lp* cell density by 48 h. Error bars are standard deviation (S.D.) for each condition, n=3. P-values are from a Student’s two-sided t-test of the difference from the monoculture (*: P<0.01, **: P<2×10-3 ). (reproduced from (*44*))

**Figure S9.**
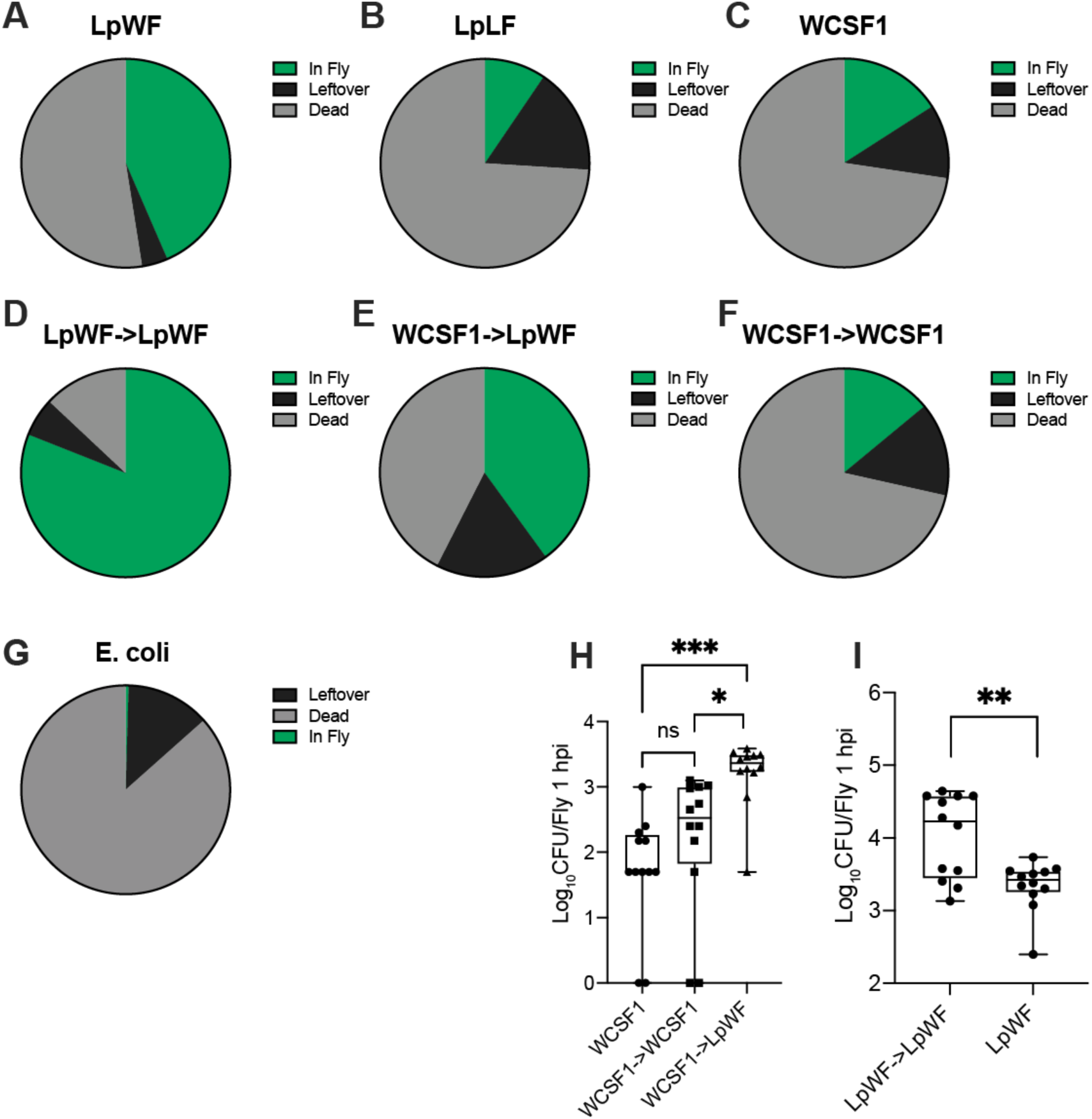
Survival and death of *Lp* strains following inoculation. A-F : The proportion of viable bacteria in the fly 1 hour post inoculation was measured alongside the bacteria remaining in the vial (leftovers), these numbers were subtracted from initial dose placed in the vial to estimate the number of bacteria killed. Proportions used for pie charts were calculated on a per fly basis. Values for flies that were fed doses of ∼10^5^ and ∼10^7^CFU/vial were combined because we did not observe significant difference (n=24 flies/bacterial strain combined from 2 vials of 12 flies/strain). The proportion of bacteria consumed (1 minus the leftover fraction) varies between strains, indicating that *LpWF* is more readily consumed by flies. These measurements were used to calculate the per-fly dose in the experiments and adjust the dose accordingly. Limit of detection = 50 CFUs. A. *LpWF* fed to germ-free flies. B. *LpLF* fed to germ-free flies. C. *WCSF1 (LpHS)* fed to germ-free flies. D. *LpWF* fed to flies pre-colonized with *LpWF*. E. *WCSF1* fed to flies pre-colonized with *LpWF*. F. *WCSF1* fed to flies pre-colonized with *WCSF1*. G. *E. coli* JM110 fed to germ-free flies. H. CFU surviving in flies fed a dose of WCSF1 (2 ×10^5^ CFU/vial or 1 x 10^4^ CFU/fly, n=12 flies). Survival of *WCSF1* after one hour was significantly higher in flies pre-colonized with *LpWF* (p=0.0006, one-way ANOVA). Survival of *WCSF1* in flies pre-colonized with *WCSF1* was not significantly higher. Survival of invading *LpWF* dose was better in flies pre-colonized with *LpWF* (4 x 10^4^ CFU/vial or 3 x 10^3^ CFU/fly, n=12 flies). p=0.0020, one-way ANOVA). I. CFU surviving in flies fed a dose of *LpWF* 2 ×10^5^ CFU/vial or 1 x 10^4^ CFU/fly, n=12 flies). Survival of *LpWF* after one hour was significantly higher in flies pre-colonized with *LpWF* (p<0.01, two-sided t-test).

---

**Movie S1.**

3-d visualization of *LpWF* and *Ai* co-colonization in the posterior proventriculus shows sectored colonization of the two strains in their respective niches. Imaging methods are the same as for Fig. 3D.

## References

1. J. J. Faith, J. L. Guruge, M. Charbonneau, S. Subramanian, H. Seedorf, A. L. Goodman, J. C. Clemente, R. Knight, A. C. Heath, R. L. Leibel, M. Rosenbaum, J. I. Gordon, The long-term stability of the human gut microbiota. Science. 341, 1237439 (2013).

2. L. A. David, A. C. Materna, J. Friedman, M. I. Campos-Baptista, M. C. Blackburn, A. Perrotta, S. E. Erdman, E. J. Alm, Host lifestyle affects human microbiota on daily timescales. Genome Biol. 15, 1–15 (2014).

3. J. G. Caporaso, C. L. Lauber, E. K. Costello, D. Berg-Lyons, A. Gonzalez, J. Stombaugh, D. Knights, P. Gajer, J. Ravel, N. Fierer, J. I. Gordon, R. Knight, Moving pictures of the human microbiome. Genome Biol. 12, R50 (2011).

4. M. C. Arrieta, L. T. Stiemsma, P. A. Dimitriu, L. Thorson, S. Russell, S. Yurist-Doutsch, B. Kuzeljevic, M. J. Gold, H. M. Britton, D. L. Lefebvre, P. Subbarao, P. Mandhane, A. Becker, K. M. McNagny, M. R. Sears, T. Kollmann, W. W. Mohn, S. E. Turvey, B. B. Finlay, Early infancy microbial and metabolic alterations affect risk of childhood asthma. Sci. Transl. Med. 7 (2015), doi:10.1126/scitranslmed.aab2271.

5. S. Subramanian, S. Huq, T. Yatsunenko, R. Haque, M. Mahfuz, M. A. Alam, A. Benezra, J. Destefano, M. F. Meier, B. D. Muegge, M. J. Barratt, L. G. VanArendonk, Q. Zhang, M. A. Province, W. A. Petri, T. Ahmed, J. I. Gordon, Persistent gut microbiota immaturity in malnourished Bangladeshi children. Nature. 510, 417–421 (2014).

6. L. A. David, C. F. Maurice, R. N. Carmody, D. B. Gootenberg, J. E. Button, B. E. Wolfe, A. V Ling, A. S. Devlin, Y. Varma, M. A. Fischbach, S. B. Biddinger, R. J. Dutton, P. J. Turnbaugh, Diet rapidly and reproducibly alters the human gut microbiome. Nature, 1–18 (2013).

7. K. M. Ng, J. A. Ferreyra, S. K. Higginbottom, J. B. Lynch, P. C. Kashyap, S. Gopinath, N. Naidu, B. Choudhury, B. C. Weimer, D. M. Monack, J. L. Sonnenburg, Microbiota-liberated host sugars facilitate post-antibiotic expansion of enteric pathogens. Nature. 502, 96–99 (2013).

8. L. Dethlefsen, D. A. Relman, Incomplete recovery and individualized responses of the human distal gut microbiota to repeated antibiotic perturbation. Proc. Natl. Acad. Sci. U. S. A. 108, 4554–4561 (2011).

9. E. D. Sonnenburg, S. A. Smits, M. Tikhonov, S. K. Higginbottom, N. S. Wingreen, J. L. Sonnenburg, Diet-induced extinctions in the gut microbiota compound over generations. Nature. 529, 212–215 (2016).

10. J. K. Kim, J. B. Lee, H. A. Jang, Y. S. Han, T. Fukatsu, B.-L. Lee, Understanding regulation of the host-mediated gut symbiont population and the symbiont-mediated host immunity in the Riptortus-Burkholderia symbiosis system. Dev. Comp. Immunol. 64, 75– 81 (2016).

11. Y. Kikuchi, T. Ohbayashi, S. Jang, P. Mergaert, Burkholderia insecticola triggers midgut closure in the bean bug Riptortus pedestris to prevent secondary bacterial infections of midgut crypts. ISME J., 1–12 (2020).

12. M. J. McFall-Ngai, The importance of microbes in animal development: lessons from the squid-vibrio symbiosis. Annu. Rev. Microbiol. 68, 177–194 (2014).

13. C. Fung, S. Tan, M. Nakajima, E. C. Skoog, L. F. Camarillo-Guerrero, J. A. Klein, T. D. Lawley, J. V. Solnick, T. Fukami, M. R. Amieva, High-resolution mapping reveals that microniches in the gastric glands control Helicobacter pylori colonization of the stomach. PLoS Biol. 17, 1–28 (2019).

14. S. M. Lee, G. P. Donaldson, Z. Mikulski, S. Boyajian, K. Ley, S. K. Mazmanian, Bacterial colonization factors control specificity and stability of the gut microbiota. Nature. 501, 426–429 (2014).

15. A. E. Douglas, The Drosophila model for microbiome research. Lab Anim. (NY). 47, 157– 164 (2018).

16. A. Saffarian, C. Mulet, B. Regnault, A. Amiot, J. Tran-Van-Nhieu, J. Ravel, I. Sobhani, P. J. Sansonetti, T. Pédron, Crypt- and Mucosa-Associated Core Microbiotas in Humans and Their Alteration in Colon Cancer Patients. MBio. 10, G351–20 (2019).

17. B. Obadia, Z. T. Güvener, V. Zhang, J. A. Ceja-Navarro, E. L. Brodie, W. W. Ja, W. B. Ludington, Probabilistic Invasion Underlies Natural Gut Microbiome Stability. Curr. Biol. 27 (2017), doi:10.1016/j.cub.2017.05.034.

18. N. A. Broderick, B. Lemaitre, Gut-associated microbes of Drosophila melanogaster. Gut Microbes. 3, 307–321 (2012).

19. C. N. A. Wong, P. Ng, A. E. Douglas, Low-diversity bacterial community in the gut of the fruitfly Drosophila melanogaster. Environ. Microbiol. 13, 1889–1900 (2011).

20. W. B. Ludington, W. W. Ja, Drosophila as a model for the gut microbiome. PLoS Pathog. 16 (2020), doi:10.1371/journal.ppat.1008398.

21. A. E. Douglas, Simple animal models for microbiome research. Nat. Rev. Microbiol., 1– 12 (2019).

22. A. W. Walters, R. C. Hughes, T. B. Call, C. J. Walker, H. Wilcox, S. C. Petersen, S. M. Rudman, P. D. Newell, A. E. Douglas, P. S. Schmidt, J. M. Chaston, The microbiota influences the Drosophila melanogasterlife history strategy. Mol. Ecol. 29, 639–653 (2020).

23. H.-Y. Lee, S.-H. Lee, J.-H. Lee, W.-J. Lee, K.-J. Min, The role of commensal microbes in the lifespan of Drosophila melanogaster. Aging (Albany. NY). 11 (2019).

24. M. A. Téfit, F. Leulier, Lactobacillus plantarumfavors the early emergence of fit and fertile adult Drosophila upon chronic undernutrition. J. Exp. Biol. 220, 900–907 (2017).

25. J. Consuegra, T. Grenier, H. Akherraz, I. Rahioui, H. Gervais, P. da Silva, F. Leulier, Metabolic cooperation among commensal bacteria supports Drosophila juvenile growth under nutritional stress. iScience, 101232 (2020).

26. S. F. Henriques, D. B. Dhakan, L. Serra, A. P. Francisco, Z. Carvalho-Santos, C. Baltazar, A. P. Elias, M. Anjos, T. Zhang, O. D. K. Maddocks, C. Ribeiro, Metabolic cross-feeding in imbalanced diets allows gut microbes to improve reproduction and alter host behaviour. Nat. Commun. 11, 4236 (2020).

27. A. L. Gould, V. Zhang, L. Lamberti, E. W. Jones, B. Obadia, N. Korasidis, A. Gavryushkin, J. M. Carlson, N. Beerenwinkel, W. B. Ludington, Microbiome interactions shape host fitness. Proc. Natl. Acad. Sci. U. S. A. 115, E11951–E11960 (2018).

28. H. Eble, M. Joswig, L. Lamberti, W. B. Ludington, Cluster partitions and fitness landscapes of the Drosophila fly microbiome. J. Math. Biol. 79 (2019), doi:10.1007/s00285-019-01381-0.

29. I. S. Pais, R. S. Valente, M. Sporniak, L. Teixeira, Drosophila melanogaster establishes a species-specific mutualistic interaction with stable gut-colonizing bacteria. PLoS Biol. 16, e2005710 (2018).

30. D. G. King, Cellular-Organization and Peritrophic Membrane Formation in the Cardia (Proventriculus) of Drosophila-Melanogaster. J. Morphol. 196, 253–282 (1988).

31. D. Tilman, Competition and Biodiversity in Spatially Structured Habitats. Ecology. 75, 2– 16 (1994).

32. K. G. Peay, M. Belisle, T. Fukami, Phylogenetic relatedness predicts priority effects in nectar yeast communities Proceedings of the Royal Society of London B: Biological Sciences. Proc. R. Soc. B Biol. Sci. (2011).

33. D. R. Amor, C. Ratzke, J. Gore, Transient invaders can induce shifts between alternative stable states of microbial communities. Sci. Adv. 6, 1–9 (2020).

34. J. Friedman, L. M. Higgins, J. Gore, Community structure follows simple assembly rules in microbial microcosms. Nat. Publ. Gr. 1, 1–7 (2017).

35. I. Martínez, M. X. Maldonado-Gomez, J. C. Gomes-Neto, Experimental evaluation of the importance of colonization history in early-life gut microbiota assembly. Elife (2018).

36. M. X. Maldonado-Gómez, I. Martínez, F. Bottacini, A. O’Callaghan, M. Ventura, D. van Sinderen, B. Hillmann, P. Vangay, D. Knights, R. W. Hutkins, J. Walter, Stable Engraftment of Bifidobacterium longum AH1206 in the Human Gut Depends on Individualized Features of the Resident Microbiome. CHOM. 20, 515–526 (2016).

37. A. L. Mattei, M. L. Riccio, F. W. Avila, M. F. Wolfner, Integrated 3D view of postmating responses by the Drosophila melanogaster female reproductive tract, obtained by micro-computed tomography scanning. Proc. Natl. Acad. Sci. 112, 8475–8480 (2015).

38. T. A. Schoborg, S. L. Smith, L. N. Smith, H. Douglas Morris, N. M. Rusan, Micro-computed tomography as a platform for exploring Drosophila development. Dev. 146 (2019), doi:10.1242/dev.176685.

39. N. Buchon, D. Osman, F. P. A. David, H. Y. Fang, J.-P. Boquete, B. Deplancke, B. Lemaitre, Morphological and molecular characterization of adult midgut compartmentalization in Drosophila. Cell Rep. 3, 1725–1738 (2013).

40. G. G. Altmann, Morphological observations on mucus-secreting nongoblet cells in the deep crypts of the rat ascending colon. Am. J. Anat. 167, 95–117 (1983).

41. A. J. Sommer, P. D. Newell, Metabolic Basis for Mutualism between Gut Bacteria and Its Impact on the Drosophila melanogaster Host. Appl. Environ. Microbiol. 85 (2019).

42. J. E. Blum, C. N. Fischer, J. Miles, J. Handelsman, Frequent Replenishment Sustains the Beneficial Microbiome of Drosophila melanogaster. MBio. 4, e00860-13-e00860-13 (2013).

43. E. S. Keebaugh, R. Yamada, B. Obadia, W. B. Ludington, W. W. Ja, Microbial Quantity Impacts Drosophila Nutrition, Development, and Lifespan. iScience. 4 (2018), doi:10.1016/j.isci.2018.06.004.

44. T. Brummel, A. Ching, L. Seroude, A. F. Simon, S. Benzer, Drosophila lifespan enhancement by exogenous bacteria. Proc. Natl. Acad. Sci. 101, 12974–12979 (2004).

45. K. Spath, S. Heinl, E. Egger, R. Grabherr, Lactobacillus plantarum and Lactobacillus buchneri as Expression Systems: Evaluation of Different Origins of Replication for the Design of Suitable Shuttle Vectors. Mol. Biotechnol. 52, 40–48 (2011).

46. C. J. Marx, M. E. Lidstrom, Development of improved versatile broad-host-range vectors for use in methylotrophs and other gram-negative bacteria. Microbiology. 147, 2065–2075 (2001).

